# Lateral Orbitofrontal Cortex Encodes Presence of Risk and Subjective Risk Preference During Decision-Making

**DOI:** 10.1101/2024.04.08.588332

**Authors:** Daniel B.K. Gabriel, Felix Havugimana, Anna E. Liley, Ivan Aguilar, Mohammed Yeasin, Nicholas W. Simon

## Abstract

Adaptive decision-making requires consideration of objective risks and rewards associated with each option, as well as subjective preference for risky/safe alternatives. Inaccurate risk/reward estimations can engender excessive risk-taking, a central trait in many psychiatric disorders. The lateral orbitofrontal cortex (lOFC) has been linked to many disorders associated with excessively risky behavior and is ideally situated to mediate risky decision-making. Here, we used single-unit electrophysiology to measure neuronal activity from lOFC of freely moving rats performing in a punishment-based risky decision-making task. Subjects chose between a small, safe reward and a large reward associated with either 0% or 50% risk of concurrent punishment. lOFC activity repeatedly encoded current risk in the environment throughout the decision-making sequence, signaling risk before, during, and after a choice. In addition, lOFC encoded reward magnitude, although this information was only evident during action selection. A Random Forest classifier successfully used neural data accurately to predict the risk of punishment in any given trial, and the ability to predict choice via lOFC activity differentiated between and risk-preferring and risk-averse rats. Finally, risk preferring subjects demonstrated reduced lOFC encoding of risk and increased encoding of reward magnitude. These findings suggest lOFC may serve as a central decision-making hub in which external, environmental information converges with internal, subjective information to guide decision-making in the face of punishment risk.

## Introduction

Healthy, effective decision-making requires accurate predictions of the risks and rewards associated with all potential options. The pursuit of rewards in the face of potentially aversive outcomes is operationalized as risky decision-making ^1–4^. While a certain degree of risk-tolerance can be beneficial, persistent choice of risky options can be maladaptive ^5–7^ and is a central facet of multiple psychopathologies, including substance use disorder (SUD), Attention-deficit/hyperactivity disorder (ADHD), and Parkinson’s disease ^1,8–14^. Thus, effective therapeutic interventions for these pathologies of decision-making requires delineating the neuronal processes that give rise to risky decisions.

The rat Risky Decision-making Task (RDT) measures preference between small, safe rewards and larger rewards accompanied by the risk of physical punishment ^3^. Critically, the RDT consistently yields a wide range of individual risk-preferences similar to that observed in human populations ^4,15–18^, including a subpopulation of “risk-preferring” rats that demonstrate preference for risky rewards regardless of high probabilities of punishment ^2,19–21^. Despite showing no distinctions from the rest of the population in pain tolerance, weight, anxiety-like behavior, or gross measures of motivation ^22^, risk-preferring rats exhibit several behavioral traits associated with vulnerability to SUD, including elevated cocaine self-administration ^21,23^, nicotine sensitivity and resilience to nicotine-evoked anxiety ^17^, sensitivity to reward-predictive cues ^24^, and increased impulsive action ^17,24,25^. Furthermore, risk-taking is associated with several neurobiological patterns including altered dopamine receptor expression in striatum and prefrontal cortex ^21,22^ and greater mesolimbic phasic dopamine release and autoreceptor function ^17,21,23,26^. Therefore, the RDT provides a method of elucidating the neural basis of individual differences in risk-taking, which may also enable understanding of a cluster of SUD-relevant traits.

Excessive risk-taking may arise from inaccurate estimation of risks or overvaluation of risk-associated rewards ^27–30^. Information about risk and reward is an element of internally generated cognitive maps, which facilitate decision-making in complex environments by providing a framework for the current state space. Lateral orbitofrontal cortex (lOFC) has been implicated in the construction and use of cognitive maps, signaling a map of state space that is accessed by other regions to track location and compare available paths within that state space^31–35^. Notably, risky decision-making is altered after disruptions of lOFC activity ^22,36–41^ and dysfunctional lOFC activity has been repeatedly observed in pathologies of decision-making like SUD ^42–49^. It is possible that lOFC contributes to excessive risk-taking by generating inaccurate representations of action-outcome contingencies ^37,50–52^. However, despite its implication in risk-taking, little is known about how lOFC processes/integrates risk and reward information within a given state space or its direct involvement with navigating between different levels of risk and reward.

Risky decision-making is not a unitary process, but rather consists of multiple stages involving distinct and overlapping cognitive processes ^6,52,53^. It is likely that critical brain regions such as lOFC transmit distinct functional signals during these different stages, highlighting the importance of experimental designs enabling segmentation of the decision-making process. Accordingly, we used a modified version of RDT that parsed decision-making into three phases: deliberation, action selection, and outcome anticipation. <u>Deliberation</u>, defined as the period before the choice is available, was isolated by training rats to perform a one second nose poke hold before the two choice options were presented. During this preparatory period, individuals generate action-outcome predictions for potential choices and evaluate the relative value of the risks and reward associated with available paths forward. This relies on internal representations of the current environmental state, as well as the probability that specific actions or cues will transition the individual from one state to another ^54–59^. After deliberation, it is necessary to commit to the chosen path and engage in <u>action selection</u>, operationalized here as the epoch immediately preceding the choice that culminates when the decision is complete. Finally, after action selection, subjects shift into <u>anticipation</u>, wherein they await impending punishments/rewards, information that is integrated into subsequent choices.

Here, we investigated the role of lOFC in punishment-based risky decision-making in adult male rats to delineate how lOFC activity encodes information on risk, reward, and choice during different stages of decision-making. First, population-level signaling was assessed by measuring changes in mean activity and selectivity across all recorded units before (deliberation), during (action selection), and after (anticipation) a decision. Then, Random Forest machine learning models^60^ were developed to assess the ability to predict both risk presence and risky/safe choice using net lOFC activity during each stage of decision-making. Next, we assessed how distinct neuronal ensembles engaged during risky decision-making encoded information about risk/safety and reward magnitude. Finally, to determine if individual differences in risk-taking are reflected in neuronal processing of risk and reward, we compared patterns of lOFC activity between “risk-preferring” and “risk-averse” rats.

## Results

### Risky decision-making

Adult male rats (n=7) were trained on a simplified version of the RDT^2,17,26,61^ designed to isolate distinct stages of decision-making: deliberation, action selection, and anticipation (Fig 1A). Rats chose between large and small magnitude reward options, with the large reward associated with either a 0% or 50% risk of foot shock punishment across two blocks of trials. As expected, the addition of risk reduced choice of the large reward (*F* (1, 5) = 8.106, *p* = .036, Fig 1B). Order of blocks (50% risk in block 1 [n=3] vs 50% risk in block 2 [n=4]) had no effect on choice behavior (*F* (1, 5) = 0.015, *p* = .906, Fig 1C). Notably, rats exhibited a range of individual differences in risk preference, such that risk of punishment elicited a 0 - 78.13% (*M* = 39.65%; Fig 1D) reduction in large reward choice.

**Figure 1.**
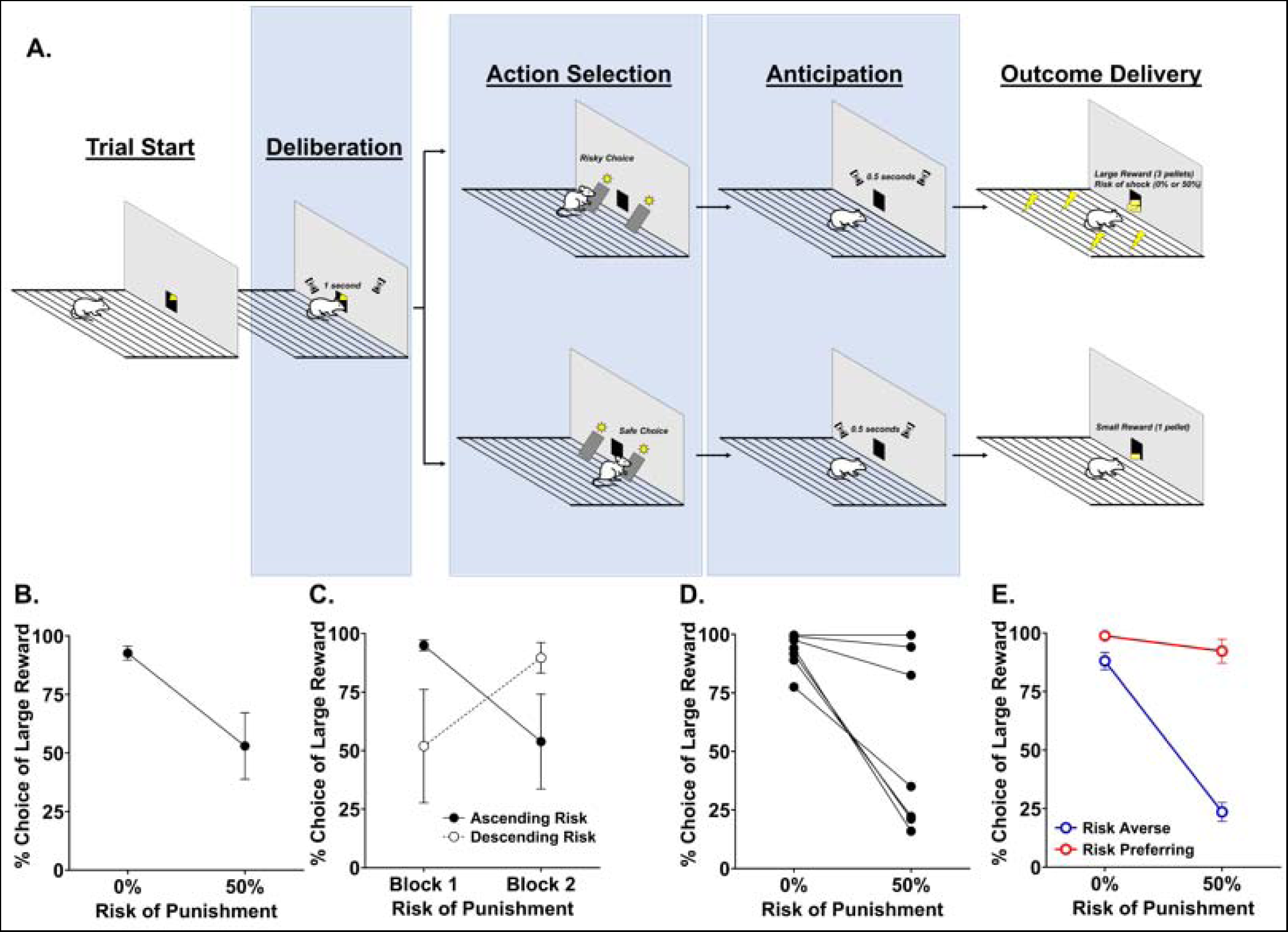
Risky Decision-Making Task (RDT) and Performance. A. At the beginning of each trial, a 1s sustained head entry into the lit trough initiates lever extension. Depression of either lever caused both levers to retract, followed by a 0.5s delay, then delivery of a small, 1 pellet reinforcer (safe choice) or a large, 3 pellet reinforcer with either a 0% or 50% risk of footshock (risky choice) depending on the current task block. Neuronal activity during was assessed during 3 distinct stages: deliberation, choice, and anticipation. B. On average, choice of large rewards decreased with a 50% risk of accompanying footshock. C. Choice of the large reward was unaffected by order the of risky (50%) vs safe (0%) blocks, i.e., ascending vs descending risk conditions. D. Rats displayed individual variability in willingness to take risks in pursuit of large rewards, and E. were grouped into risk averse and risk preferring groups. Data displayed in B,C, and E depict mean ± SEM.

Latency to select large rewards was unaffected by the presence of risk (*F* (1, 6) = 1.541, *p =* .260) and choice latency did not differ between large and small rewards within the risky block (*F* (1,6) = .001, *p* = .980). Incomplete trials (*M* = 134.08 ± 24.67) due to either omitted responses or failed initiation holds were repeated until completion during each session of RDT. Risk preference was not correlated with the number of repeated trials (*r* (6) = -.199, *p =* .668). Incomplete trials did not differ between 0% and 50% risk conditions (*F* (1,6) < .000, *p* = .990) and were unaffected by the order of risk conditions (*F* (1,6) = 1.244, *p* = .315). Furthermore, incomplete trials occurred equally due to failed trial initiations and omitted responses (*t* (6) = 1.118, *p =* .306) and risk of punishment did not alter either initiation failure (*F* (1,6) = 0.496, *p* = .508) or trial omission (*F* (1,6) = 0.031, *p* = .865). Collectively, these data suggest that the presence of risk did not alter overall task engagement/performance.

### LOFC activity is modulated by multiple events during risky decision-making

Neuronal activity was recorded through 16 channel drivable microwire electrode arrays implanted in lOFC (Fig 2A) over 36 sessions of RDT. 554 single units were sorted based on spike waveform topology (Fig 2B) and PCA clustering (Fig 2C) and event-evoked changes in neuronal activity was analyzed during 3 distinct task epochs: deliberation, action selection, and outcome anticipation (Supplemental Table 1). Based on our selectivity criteria, 78% of recorded cells (433 units) were either activated or suppressed during one or more of these epochs (Fig 2D-F), demonstrating that lOFC neurons are bidirectionally engaged by different events throughout risky decision-making.

**Figure 2.**
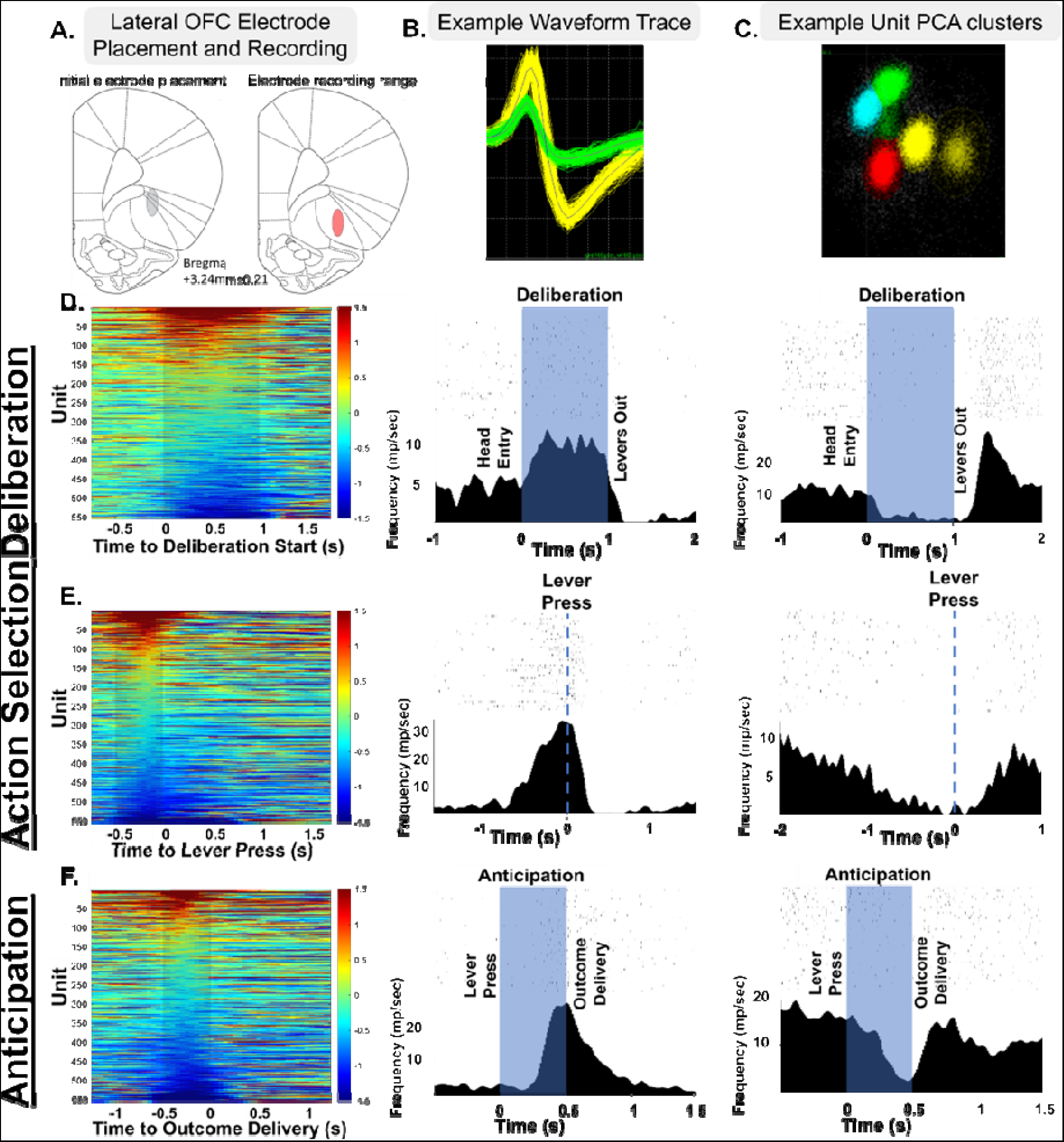
Population lOFC activity is modulated by different stages of risky decision-making. **A.** Electrodes were positioned in the lateral orbitofrontal cortex. **B.** Recorded spike waveforms were sorted into individual neuronal units based on waveform topology. **C.** Principal Component Analysis (PCA) was us d to generate principal component (PC) loadings for individual spikes and confirm accuracy of single-unit sorting via PC clustering. **D,E,F.** Heat plots of individual unit activity during risky decision-making deliberation (**D.**), choice (**E.**), and anticipation (**F.**), reveal that activity is bidirectionally modulated by each event. Units were sorted in order of average firing rate during the event. Next to each heat plot, raster plots show representative examples of individual units that were phasically activated (col 2) or suppressed (col 3) during risky decision-making. Mean neuronal spiking is displayed as a histogram in each raster plot.

### LOFC population activity encodes punishment risk during multiple events throughout decision-making

First, we compared population activity between the risk-free and 50% risk of punishment blocks during different stages of RDT (Fig 3A,E,I). During deliberation, lOFC population activity was elevated in the presence of risk (*F* (1, 552) = 4.575, *p* = .033, Fig 3C) and 52.5% of activated or suppressed units changed selectivity between the 0% and 50% risk conditions. During action selection, there was a significant risk condition x bin interaction (*F* (4, 2208) = 10.001, *p* < .001, Fig 3G), such that the presence of risk evoked a shift in firing rates from being suppressed to activated, and 51.1% of units changed selective responses between risk conditions. Finally, during anticipation there was a significant risk x bin interaction (*F* (9, 4968) = 1.915, *p* = .045, Fig 3K), such that population firing rate was again shifted from being suppressed to activated in the risky condition, and 65.8% of units changed their selectivity between blocks. Therefore, the presence of risk is encoded by an increase in lOFC population activity that manifests repeatedly throughout decision-making.

**Figure 3.**
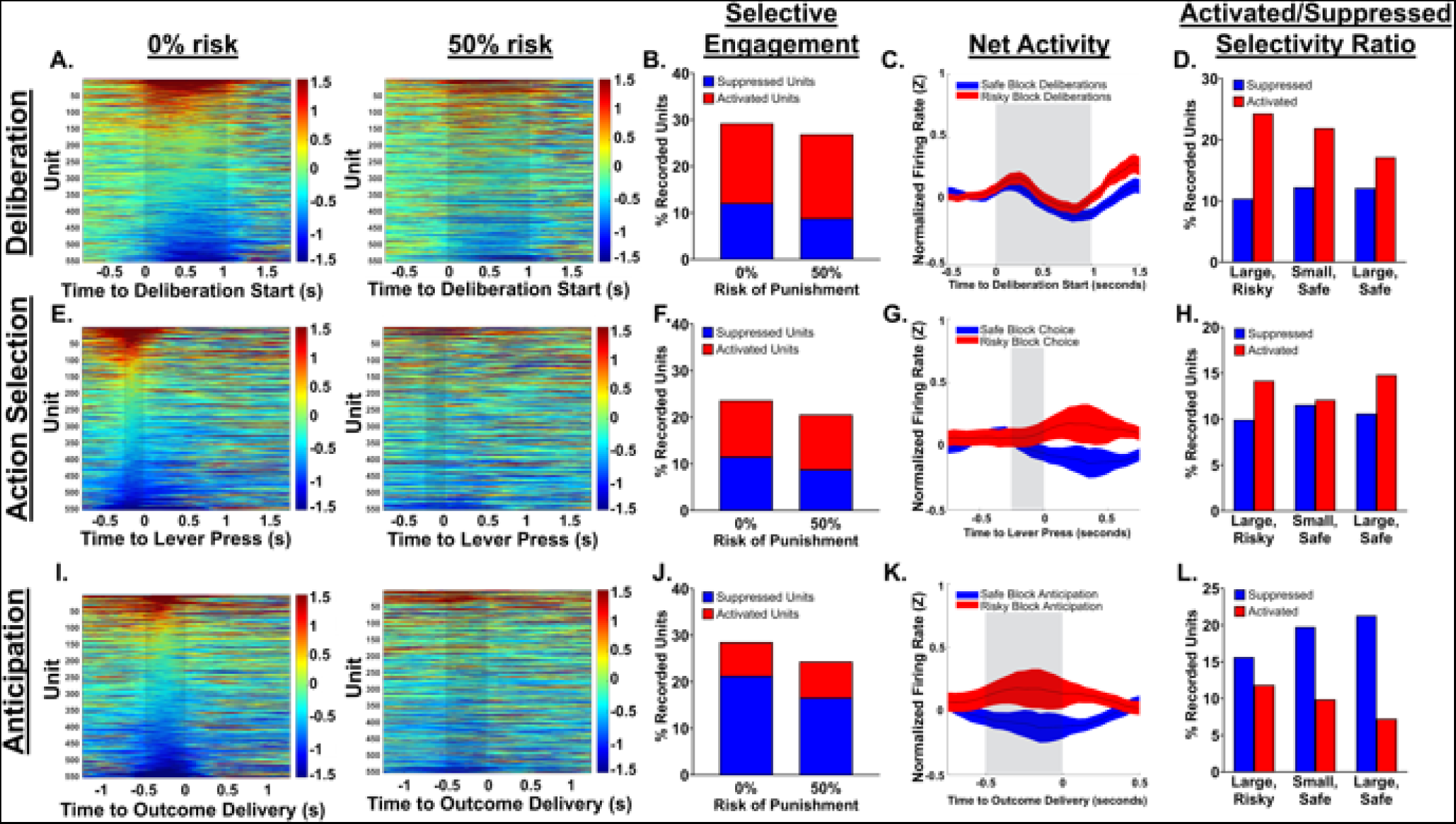
The presence of risk alters population lOFC signaling throughout the risky decision-making process. **A,E,I.** Heat plots showing activity of individual units throughout risky decision-making deliberation (**A.**), action selection (**E.**), and anticipation (**I.**). Heat plots of the same units 50% risky block reveal changes in evoked single unit activity with the addition of risk. Within each heat plot, rows are sorted based on average firing rate in the 0% risk block. **B,F,J.** Neuronal engagement (%) of units with event-evoked activation or suppression) did not differ between risk conditions. **C,G,K.** Population firing rate of all recorded units across the 0%/50% risk conditions. Net activity is consistent between risk conditions during the first half of deliberation (**C.**) but begins to increase in the risky block as deliberation nears its end. Net activity fully diverges at the moment of choice (**G.**), decreasing in the 0% risk condition and increasing in the 50% risk condition. When risk is present in the environment, net activity is elevated during anticipation (**K.**) of chosen rewards. **D,H,L.** Trials containing large, risky rewards vs large, safe rewards are encoded via an increased ratio of selective activations to suppressions during deliberation (**D.**) and a decreased ratio during anticipation (**L.**). This selectivity ratio did not differentiate between trials during action selection (**H.**), nor between trials resulting in small vs large rewards regardless of risk level. Data depicted in C,G, and K represent mean ±SEM.

Overall lOFC engagement (total units either activated or suppressed by an event, Supplemental Table 1) was comparable between risk and risk-free blocks (Fig 3B,F,J). To assess risk-evoked changes in activated vs suppressed selective signaling, we measured how the ratio of activated to suppressed units was altered during each stage of RDT. During deliberation, presence of risk was encoded as an increase in the proportion of activated to suppressed units (χ^2^ (1, n = 326) = 4.686, *p* = .030, Fig 3D), whereas anticipation of large, risky rewards was associated with an increase in suppressed to activated units ( χ^2^ (1, n = 288) = 10.125, *p* = .001, Fig 3L). The ratio of selective activations/suppressions did not differ between risk conditions during action selection ( χ^2^ (1, n = 245) = 1.431, *p* = .232, Fig 3H). Critically, in trials where rats chose small safe rewards this activated/suppressed ratio did not differ from large, safe reward choice during deliberation, action selection, or anticipation (all χ^2^ < 2.6, *p*-values > .1), suggesting that changes in population dynamics observed during risky choice were not related to reward magnitude or the overall value of a choice. In summary, the presence of risk is encoded by bidirectional shifts in the ratio of activated:suppressed units during the deliberation before a choice and outcome anticipation after a choice, but not during action selection.

### Using Machine Learning to Decode Net lOFC Activity

We next trained a Random Forest classifier using firing rates of all recorded units to decode lOFC signaling of current risk associated with the large reward on a trial-by-trial basis. We found lOFC population activity accurately predicts the current risk of punishment associated with large rewards, with an average accuracy of 75% ± 2.65 across all three stages of decision-making (Fig 4A). Interestingly, the model was more accurate during deliberation and anticipation than during action selection. Measures of precision, recall, and F1 score were comparable to accuracy. In summary, net lOFC activity can be used to predict presence/absence of risk with high accuracy throughout all stages of decision-making, with the highest accuracy occurring during deliberation.

**Figure 4.**
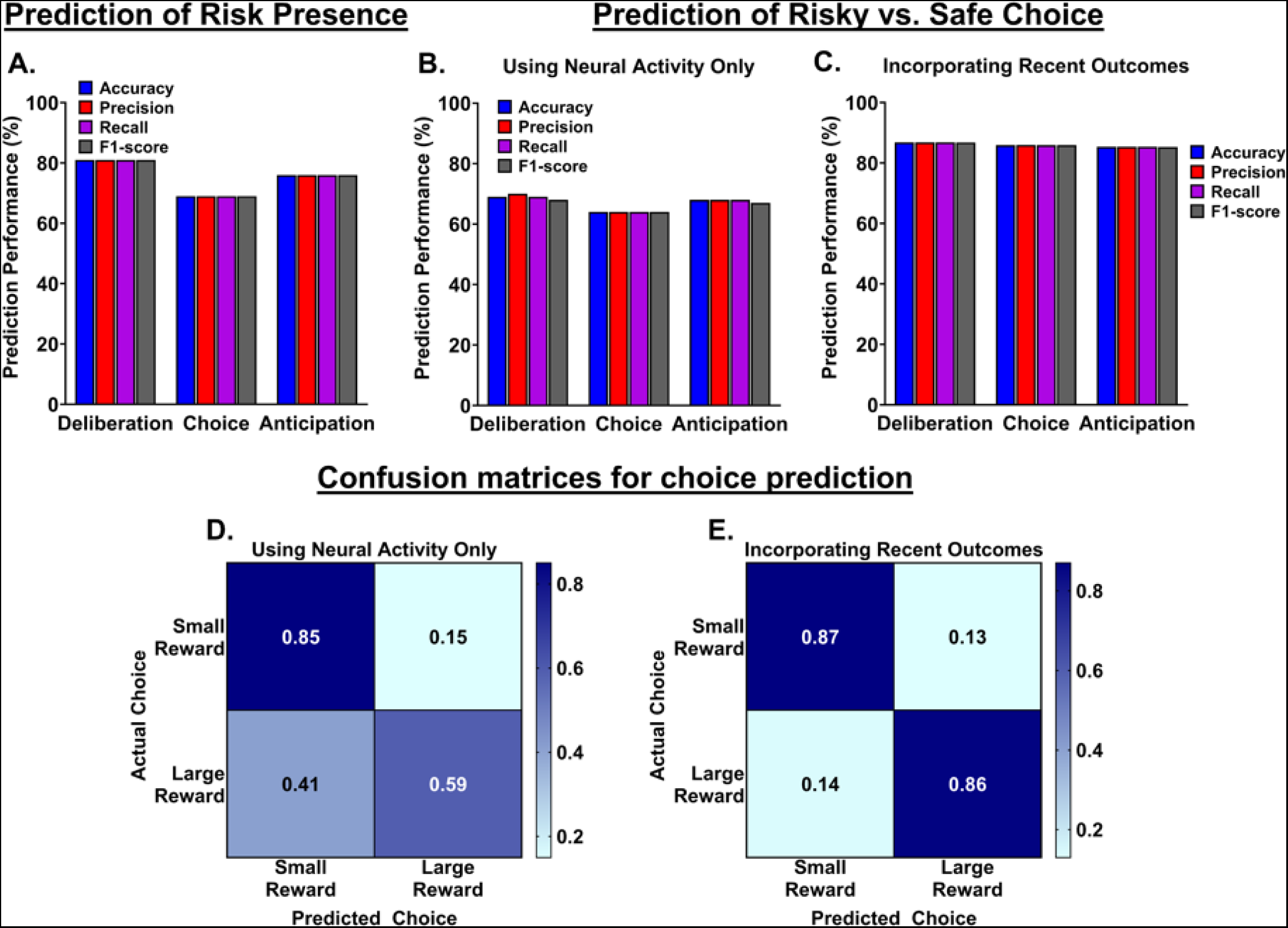
Populations activity decoding to predict current risk conditions and choice on a trial-by-trial basis. **A.** A parameter-optimized Random Forest (RF) model accurately predicts whether any given trial occurs in the risky or safe task block using population activity. **B.** Trial-by-trial choice is predicted above chance using population activity during different events in decision-making. **C.** Choice prediction improves when previous trial outcome is incorporated into the RF classifier’s dataset. **D.** Average confusion matrix for choice prediction model using only neural data, averaged across all three trial stages. Misclassification of large, risky choice trials as small, safe choice trials accounted for the majority of model errors. **E.** Average confusion matrix for choice prediction model incorporating recent events with neural data. When provided previous trial outcomes as a categorical variable to contextualize neuronal activity, model performance drastically increased, driven by improved classification of large, risky choices. Data are displayed as proportion of choices of each type predicted correctly or incorrectly; for example, .85 of small reward choice trials and .59 of large reward trials were predicted accurately in D.

The previous data suggest a role for lOFC in signaling risk throughout decision-making, though it remains unclear if lOFC actively guides behavior through the risky decision-making trial. To explore the possibility of a more direct role for lOFC in guiding choice behavior, we trained a Random Forest classifier on population activity in the risky block to predict whether a given trial resulted in large risky or small safe reward choice. The model correctly identified whether 66.67% ± 3.67 of trials resulted in a risky or safe choice (Fig 4B), averaged across the 3 trial epochs. Though this resulted in a model exceeded chance levels, it was an underperformance compared to the previous model trained to predict the presence of risk (Fig 4A).

A commonly proposed role for lOFC is to inform evaluative processes by incorporating recent events with existing associative models^62–65^. Accordingly, we retrained the model on the same neuronal data set with the addition of a categorical variable indicating the outcome of the previous choice trial (small reward, large reward no footshock, or large reward + footshock). By incorporating recent events, the Random Forest classifier improved across all three stages of decision-making, reaching an average classification accuracy of 86.01% ± 0.42 (Fig 4C). Confusion matrices revealed that this improved performance was primarily driven by better classification of large, risky choices. When trained on neural data alone, 41% of trials containing large, risky choice were misclassified as small, safe choice trials (Fig 4D). Incorporating previous trial outcomes with the model reduced misclassification rate to 14% (Fig 4E). This suggests that when subjects engage in risky behaviors, lOFC (or a downstream region) incorporates the previous outcome into the evaluative process. Conversely, misclassification of small, safe choice trials was consistently low (15% ➔ 13%), indicating that the previous outcome is only informative after decisions with an element of risk. Thus, lOFC direction of risky behavior may involve integration of the outcome of the previous risky decision.

### Subgroups of LOFC neurons encode presence of risk regardless of choice

To investigate lOFC processing of risky decision-making on a more granular level, we analyzed activity within subpopulations of units phasically activated or suppressed during different task events. Separate neuronal subpopulations displayed suppressed activity during deliberation (n=67), action selection (n=64), and anticipation (n=118) in the absence of risk. However, when risk of punishment was present in the task space, this suppression was significantly reduced (deliberation: *F* (2,162) = 11.415, *p* < .001; action selection: *F* (2,165) = 18.686, *p* < .001; anticipation: *F* (2,307) = 41.592, *p* < .001; Fig 5A,D,G). Interestingly, this risk-mediated attenuation of suppression occurred regardless of whether subjects chose the large, risky (Fig 5B,E,H) or small, safe (Fig 5C,F,I) rewards (pairwise comparisons: all *p*-values < .05). Thus, lOFC encode presence/absence of risk rather than comparing risk level between concurrently available paths through the task space (risky vs safe choice).

**Figure 5.**
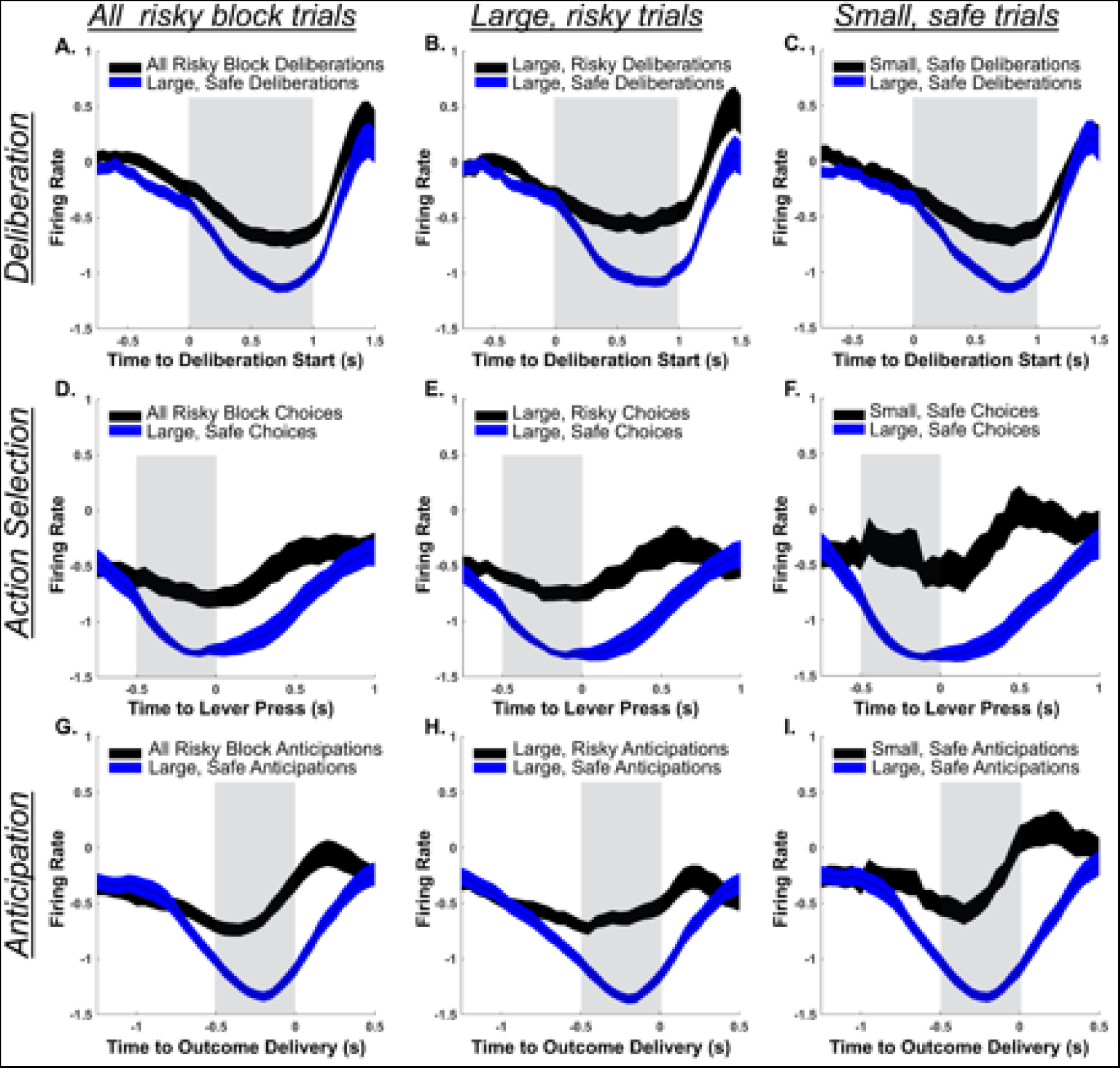
lOFC signals the presence of risk throughout decision-making. Distinct neuronal subpopulations were phasically suppressed during deliberation (row 1), action selection (row 2), and anticipation (row 3) in the 0% risk block of trials. A,D,G. This selective suppression was attenuated during the 50% risk block across all events. Notably, this attenuated response occurred regardless of whether subjects chose large, risky (B,E,H.) or small, safe (C,F,I.) reward, suggesting it is signaling changes in environmental risk rather than encoding risky vs safe choice. All data are depicted as mean±SEM.

### Selectively activated lOFC units encode reward magnitude during action selection

Having identified evidence that lOFC neuronal subpopulations encode risk, we next examined reward magnitude, a well-known feature of lOFC signaling ^66–68^. We identified 111 units selectively activated during action selection of either large, safe or small, safe rewards. These subpopulations were classified as either “large magnitude-encoding” or “small magnitude-encoding” based on the reward choice (large, safe or small, safe) during which they exhibited the highest degree of phasic activation. There was no significant difference in peak firing rate between the two magnitude-encoding subpopulations (*F* (1,129) = 0.042, *p* = .839). Critically, each of these subpopulations discriminated between small and large rewards via alterations in phasic activation. Large magnitude-encoding neurons selectively activated more during choice of large (risky or safe) than small rewards (*F* (2,177) = 5.609, *p* = .004), with individual comparisons confirming comparable activity between large risky and large safe reward choices choice (*p* = .946, Fig 6A) and a robust reduction in firing rate for small reward compared to large, safe reward choice (*p* = .003, Fig 6B). Similarly, small magnitude-encoding neurons exhibited greater neuronal activity during small reward choice (*F* (2,178) = 7.202, *p* < .001) compared to either large safe (*p* < .001, Fig 6C) or large risky (*p* = .025, Fig 6D) reward choice. Thus, separate subpopulations of neurons in lOFC encode reward magnitude during action selection separately from the presence of risk.

**Figure 6.**
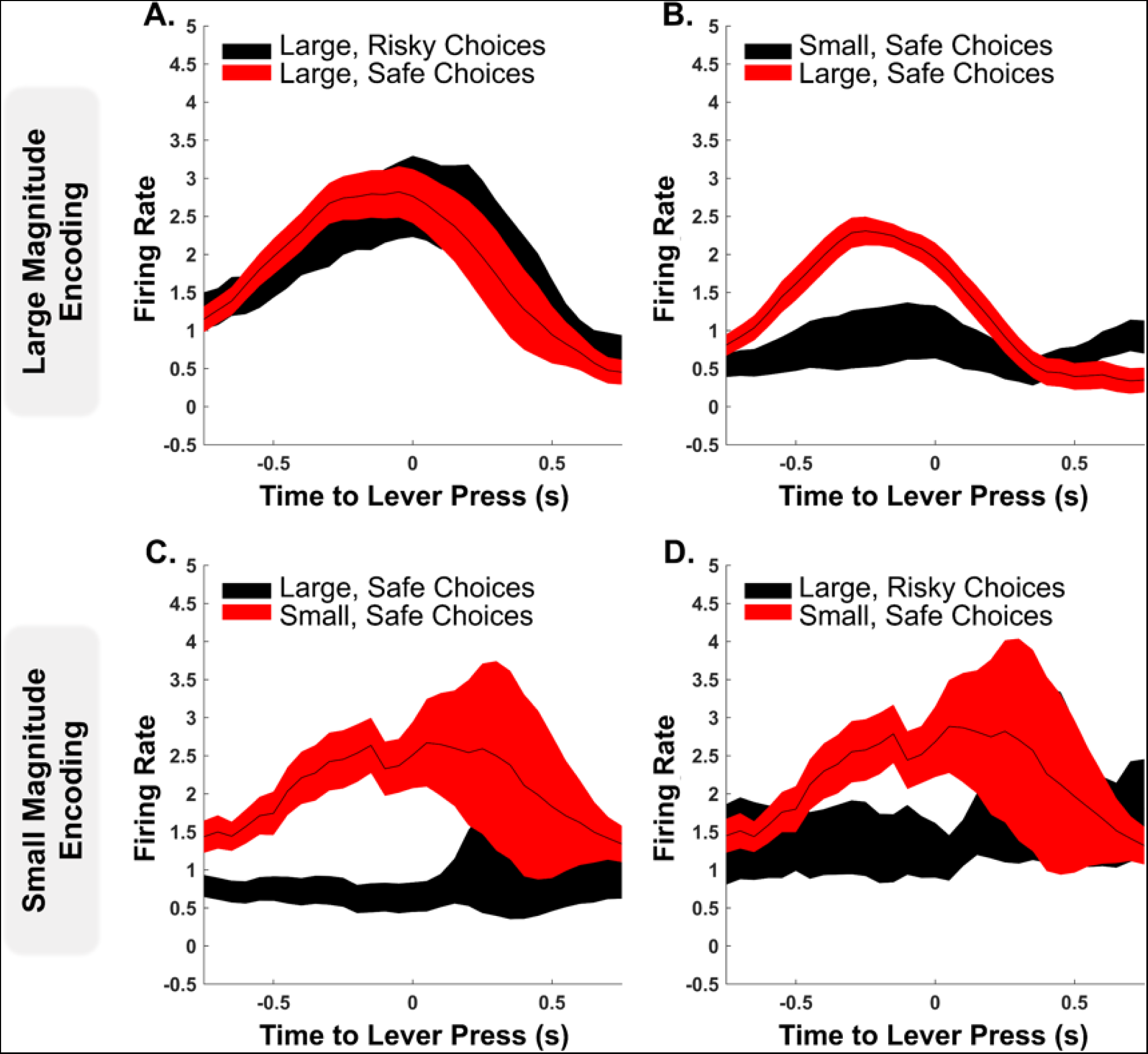
lOFC encodes reward magnitude during action selection regardless of risk. **A,B.** Units selectively activated during choice of large, safe rewards in the safe block exhibited similar activations in the risky block during large, risky choice (**A.**), but not during small, safe choice (**B.**). **C,D.** Units selectively activated during choice of small, safe rewards in the risky block were not activated when choosing either large, safe (**C.**) or large, risky (D.) rewards. All data depicted as mean±SEM.

### LOFC activity underlying individual differences in risk-taking

#### Individual differences in risk-taking are associated with differences in lOFC risk processing

Rats were grouped as either risk-preferring or risk-averse based on preference for risky vs safe choice (Fig 1E). To determine if individual differences in risk-taking were mirrored by differences in lOFC activity, we compared the processing of both risk and reward magnitude between these groups. We identified a subset of neurons which were suppressed during deliberation prior to risky choice in both risk-averse (units=39) and risk-preferring (units=10) rats. This neuronal subpopulation showed divergent activity in response to risky and safe rewards as a function of individual risk preference (*F* (1,47) = 11.859, *p* = .001). In risk-averse rats, this suppression was significant attenuated during deliberation prior to large, safe choice (*p* < .001, Fig 7A). However, in risk-preferring rats, this subpopulation exhibited nearly identical suppression between deliberations preceding large, risky and large, safe choice (*p* = .670, Fig 7B). A similar dichotomy in risk encoding between risk preferring and averse rats occurred during outcome anticipation (*F* (1,77) = 5.326, *p* = .024). In both groups, distinct neural subpopulations exhibited selective suppressions during anticipation of risky rewards. In risk-averse rats, this selective suppression was absent during anticipation of large, safe rewards (n=45, *p* < .001, Fig 7C). Conversely, in risk-preferring rats, this subpopulation did not discriminate between large, safe and large, risky choice (n=29, *p* = .614, Fig 7D). Thus, lOFC encoding of risk differs based on individual differences in risk-preference, with risk-preferring rats showing reduced encoding of risk during both prior to (deliberation) and following (outcome anticipation) a choice.

**Figure 7.**
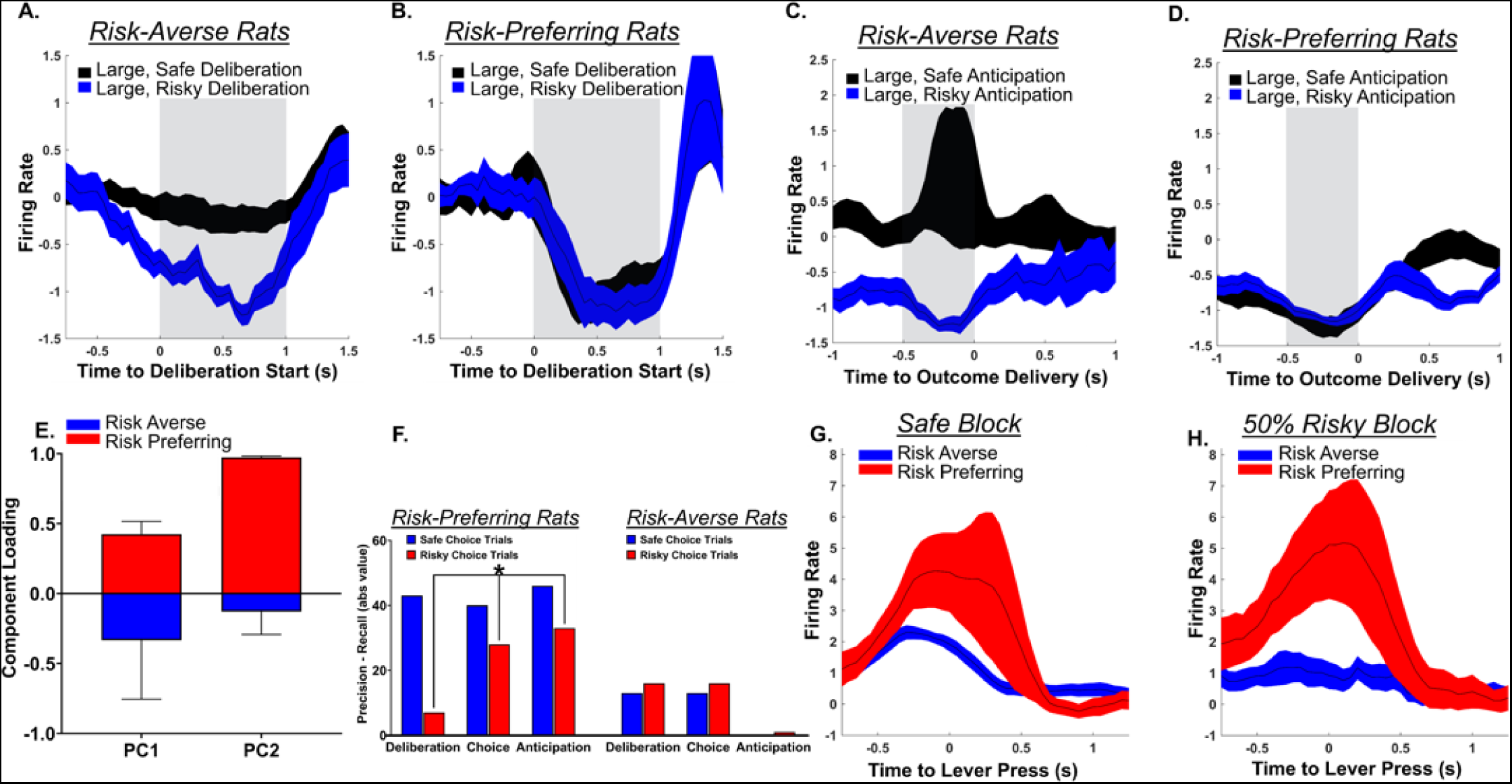
Individual differences in risk-taking are associated with differences in lOFC risk processing and increased lOFC reward sensitivity. A,B. In both risk-averse and -preferring rats, distinct neuronal subpopulations were suppressed during deliberations preceding large, risky choice. In risk-averse rats (A.), this suppression is absent during deliberations preceding large, safe choices. However, risk-preferring rats (B.) exhibit identical suppressions during deliberation prior to both safe and risky large reward choice, indicating a lack of neuronal sensitivity to risk. C,D. Similarly, selective subpopulations are suppressed during anticipation of large, risky rewards in both risk-averse and -preferring rats. In risk-averse rats (C.), this suppression is absent during anticipation of large, safe rewards, while risk-preferring rats (D.) demonstrate identical suppression during anticipation of large safe and risky rewards. E. Principal components 1 and 2 neatly distinguished between risk-preferring and risk-averse rats. Performance metrics component loadings for classification of risky and safe choice trials correlated with subjective risk preference. F. In risk-preferring rats, but not risk-averse, precision and recall diverged on trials containing choice of non-preferred rewards. G. Neurons activated during action selection of large, safe rewards exhibit stronger activation in risk-preferring rats than risk-averse rats. H. When large rewards include a 50% risk of shock, this activation is maintained in risk-preferring rats but muted in risk-averse rats. Data in A-D & G-H depicted as mean±SEM.

We trained a Random Forest classifier to further explore how individual risk preferences are encoded by lOFC signaling on a trial-by-trial basis. This model matched risk preference with population activity to predict 1) current risk associated with large rewards, and 2) choice between large risky vs small safe rewards. F1-scores — the harmonic mean of precision and recall, and a normalized metric of model performance^69^ — were used to compare model performance between risk groups and between safe and risky choice). In both risk-preferring and -averse rats, lOFC activity encoded presence/absence of risk associated with large rewards (Supplemental 2A,B). OFC encoding of risk condition was consistent (<5% variance between metrics) at each trial stage for both risk-preferring and -averse rats (Supplemental 2A,B) and there were no strong differences in F1-score (>5%) between risk groups (Supplemental 2C). This lack of group difference suggests that subjective preference for/against large, risky rewards does not originate from deficient LOFC prediction of the presence of risk.

However, while lOFC activity reflected the presence/absence of risk comparably between risk groups, differences emerged in LOFC encoding of risky vs. safe choice. Using linear regression, we determined that Random Forest performance metrics predicted choice of risky vs safe rewards (*r^2^*= 0.627, *F* (3,11) = 4.482, *p* = .040). We then used principal components analysis to assess how lOFC encoding varies based on subjective risk preference. Performance metrics loaded on the first and second components along lines according to risk-group (Fig 7E), indicating that variance in lOFC encoding of decision-making largely correlates with subjective risk preference. Finally, a logistic regression determined that decoded lOFC signaling of risky vs safe choices predicted subjective risk preference (χ^2^ (6, N =36) = 16.636, *p* = .011). <u>This indicates that lOFC encoding of choice is linked to subjective preference for risky vs safe rewards.</u>

In risk-preferring rats, precision, a measure of model classification, was elevated (>5%) above other performance metrics during choice and anticipation (Supplemental 2D), whereas in risk-averse rats, it was elevated during deliberation and choice (Supplemental 2E). These group differences in encoding risky vs safe choice were further exemplified by differences in F1-score during deliberation and choice (Supplemental 2F). Within each risk group, differences in precision— a measure of false positives— and recall— a measure of false negatives— were further compared at the level of discrete classes (safe choice and risky choice, Supplemental 2G,H). Separate Repeated Measures ANOVA (class x metric) revealed that precision and recall metrics of lOFC encoding were significantly lower when classifying non-preferred rewards in both risk-preferring (*F* (1,2) = 131.942, *p* = .007, Supplemental 2G_solid bars_) and risk-averse (*F* (1,2) = 85.563, *p* = .011, Supplemental 2H_slashed bars_) rats. Furthermore, risk-preferring rats exhibited a class x metric interaction (*F* (1,2) = 60.356, *p* = .016) in the form of a divergence between precision and recall when classifying safe choice trials. Risk-preferring rats’ average precision across each trial stage fell 43% ± 1.73 below recall during safe choice trials, but not during risky choice trials (pairwise *p*-values: safe class = .002; risky class = .104, Fig 7F_left_). This reflects an increase in the rate of false positives in relation to the rate of false negatives when identifying safe choice trials, whereas precision and recall were comparable when classifying risky choice trials. <u>Thus, despite accurate LOFC encoding of risk vs safety in the environment, risk-preferring rats appear to lack a distinct LOFC state associated with safe behavior, especially during action selection and anticipation, wherein precision fell below 10%.</u> This divergence between precision and recall did not occur for risk-averse rats (*F* (1,2) = 3.610, *p* = .198, Fig 7F_right_).

### Risk-taking is linked to increased lOFC reward sensitivity

To test if excessive risk-taking may arise from biased signaling of large vs small rewards, we examined how reward magnitude discrimination in lOFC differed based on individual differences in risk-taking. In risk-preferring rats, large reward magnitude-encoding neurons exhibited a greater phasic increase in firing rate during action selection than in risk-averse rats when choosing large rewards (*F* (1,174) = 4.913, *p* = .028; large, safe: *p* = .027, Fig 7G; large, risky: *p* = .004, Fig 7H), but did not differ between risk groups when choosing small, safe rewards (*p* = .612). This suggests that risk-taking may be caused in part by elevated reward magnitude signals during action selection.

## Discussion

Decision-making is a complex cognitive process that requires accurate evaluation of internal and external factors, including assessment of the risk of punishment and reward magnitude associated with all options within the current environmental state ^2,4,70,71^. We found that lOFC encodes the presence of risk throughout decision-making, with both population activity and selectively engaged neuronal subpopulations signaling impending risk during deliberation prior to a choice, action selection, and post-choice outcome anticipation. Moreover, lOFC activity can be decoded to accurately predict current risk levels as well as risky vs safe choice on a trial-by-trial basis. LOFC also encodes reward magnitude, although this pattern of activity was only evident during action selection. Finally, both reward and risk encoding differed based on subjective risk preference, with risk-preferring rats exhibiting attenuated lOFC discrimination between risky and safe conditions, elevated lOFC activation during large rewards, and reduced ability to predict safe choice via lOFC activity.

### lOFC encodes risk of punishment throughout decision-making

Neuronal activity in OFC encodes appetitive and aversive valence ^72–74^, and appetitive and aversive information converges at the level of individual neurons in OFC ^75–78^. Moreover, single-units in lOFC adaptively encode changes in information about available options, updating activity to match behavior with available reward information ^79–82^. This dynamic encoding of motivational stimuli in the environment dovetails with the lOFC discrimination between risky vs non-risky conditions observed here. It is important to note that lOFC encoded risk not only during expectation of risky outcomes, but also during pre-choice deliberation and the choice itself. Critically, lOFC risk encoding was not linked to decision-making on a trial-by-trial basis, with risk-related activity occurring comparably regardless of engagement in large, risky or small, safe choice.

This repetitive risk-encoding throughout the decision-making process, manifesting as reduced suppression of lOFC activity, has several potential explanations. One explanation is the production of a robust signal that is more resistant to degradation ^83^, ensuring that highly salient punishment risk is properly incorporated into the decision-making process. Another possibility is that lOFC may subserve a different function at each stage of decision-making, which is supported by the variation in predictive accuracy achieved by Random Forests decoding classifier across trial stages (Fig 4A). During deliberation, risk signals may contribute to evaluation of risk prior to action selection. During choice itself, reduced suppression of lOFC activity in the presence of risk may reflect increased output to regions associated with either invigorating or suppressing motivated action. Finally, during anticipation lOFC signals may facilitate activation of downstream regions associated with threat detection or reward seeking/consumption. This multifunctional role for lOFC in the decision-making process is supported by confusion matrices at each stage of decision-making. During deliberation, the Random Forest model misclassified extremely few trials as being in the risky vs safe task conditions (Supplemental 3A). However, misclassification became more common during action selection (Supplemental 3B) and anticipation (Supplemental 3C) and occurred with similar frequency during these latter stages of the decision-making process. This suggests that lOFC activity may reflect changes in cognitive requirements as decision-making progresses from evaluative to invigorating to anticipatory. Further research is needed to investigate this possibility and identify the brain regions influenced by these risk signals, as well as whether risk encoding arises from inhibitory or excitatory neurons.

### lOFC encoding of risky choice

lOFC activity predicted impending risky choices via changes in ratio of activated to suppressed neurons during pre-choice deliberation. This signal may directly reflect the neuronal state in which subjects are sufficiently motivated to engage in risk-taking, as enhanced OFC activity plays a causal role in willingness to endure punishment in pursuit of reward ^84^, and OFC signaling has been repeatedly linked to punishment sensitivity, though in inconsistent directions ^48,85,86^. This activation/suppression ratio may also reflect a model-based value estimate in which value of the large reward is incorporated with value of potential rewards in future states and the probability of transitioning into those states ^58,87^. Critically, it is unlikely that the increased number of activated units preceding risky choice was related to reward magnitude, as activated-suppressed ratio was comparable between expectation of small and large safe rewards. The ratio of activated to suppressed units was also altered during outcome anticipation following risky vs safe choice. This could reflect changes in lOFC communication with regions associated with dread or consolidated reward value ^66,81,91^, as lOFC projects information about impending outcomes to basolateral amygdala, nucleus accumbens, and dorsal striatum^92–94^. In summary, in addition to processing information about risk in the environment, lOFC encodes willingness to pursue a risky outcome both before and after choice. It is possible that these signals may be necessary to invigorate reward seeking in the presence of threat.

### Reward magnitude is encoded within lOFC during action selection

During action selection, distinct lOFC neuronal subpopulations encoded reward magnitude separately from risk. Event-selective ensembles discriminated between different outcomes by activating during either large or small reward choice (Fig 6), then this phasic activation was attenuated during choice of the other magnitude. These magnitude encoding subpopulations did not discriminate between risky and non-risky large rewards, suggesting that these signals are specific to reward size without integrating the presence of risk into the expected outcome. This aligns with previous research indicating that lOFC encodes both value and reward magnitude associated with specific actions ^34,66,68,95,96^, and encodes magnitude separately from subjective value assigned to a given reward ^66,82,92^. Additionally, the encoding of reward magnitude via phasic lOFC activation has been previously observed ^92,97–99^. Notably, while ensembles encoded the presence of risk both before and after a choice, reinforcer magnitude was only signaled during action selection.

### lOFC encoding of risk and reward varies with individual differences in subjective risk preference

The abridged version of RDT used here produced robust individual differences in risk preference, replicating previous observations with standard RDT ^3,4,17–19,22,23,25,26,100^. Differences in subjective risk preference were accompanied by differences in risk and reward magnitude signaling in lOFC, and Random Forest choice classification using trial X trial neuronal data was sufficient to predict subjective risk preference. Risk-averse rats exhibited selectively suppressed neuronal subpopulations during deliberation preceding large risky reward choice, but this signal did not precede large safe choice. In contrast, risk-preferring rats displayed an identical suppression in activity between risky and safe large rewards. This may reflect reduced lOFC processing of impending risk in risk-preferring rats, resulting in failure to properly integrate risk into mental representations of state space during evaluation. Alternatively, this attenuated risk encoding may reflect risk-takers attributing less salience to impending punishment, driving increased tolerance of risky outcomes. It is unlikely that reduced risk salience is directly related to physical sensitivity to punishment, as subjective risk preference in RDT is not associated with shock sensitivity or general pain tolerance, and is not influenced by acute exposure to an opioid analgesic ^22,100^. Nor is it likely that increased risk-taking arises from a lack of awareness that risk is present, as there are no substantial differences in block-prediction performance metrics (Supplemental 2A-C) or confusion matrices (Supplemental 3D-I) between risk-preferring and risk-averse rats.

A predisposition toward excessive risk-taking may arise from elevated lOFC processing of large rewards, possibly driving increased willingness to seek these rewards despite the presence of risk. While large reward-encoding neuronal subpopulations were identified within both risk groups, these neurons generated greater phasic activation in risk-preferring rats. Additionally, while risk-averse rats exhibited a blunted signal from these large reward-encoding units in the presence of risk, risk-preferring rats exhibited consistent signal strength when choosing large rewards regardless of the associated risk. This aligns with another report that elevated risk-preference is associated with increased lOFC processing of reward magnitude ^101^. Collectively, these data suggest that excessive risk-taking may arise from both increased reward-evoked signaling and reduced risk-signaling in lOFC, resulting in a bias toward risky paths through the state space.

It is also possible that risk-preferring rats’ persistent choice of risky rewards is caused by elevated habit formation, leading to failure to adapt to changes in action-outcome contingencies within a session. However, all rats here had comparable levels of training, and individual risk preference is uncorrelated with susceptibility to habit formation ^17^. Additionally, by requiring subjects to perform a sustained nose poke hold to initiate lever extension at the start of each trial, the current design imposes a break in the standard stimulus-response pattern underlying habitual behavior. This pause may promote goal-directed behavior by forcing a period for evaluation of outcomes associated with upcoming options. Finally, a subset of subjects performed the task with the risky block preceding the safe block, suggesting that persistent risky choice cannot solely be explained by inflexible choice of the previously risk-free large reward.

### Risky Decision-making and Cognitive Mapping

Cognitive maps facilitate rational behavior by providing 1.) a mental model of the states comprising the current decision environment, and 2.) the rules and probabilities associated with transitioning between distinct states ^55,56,58,87^. Neuronal activity in lOFC contributes to tracking goal-directed behavior ^66,102–105^ and mediates the associative learning used to guide action-selection in downstream regions ^43,78,106–112^. Thus, while lOFC is not proposed to directly invigorate choice ^113–116^, it is critical to the construction, updating, and use of cognitive maps ^31,32,56,99,117^. Here, we identified an active role for lOFC in generating dynamic representations of state space by encoding both external (risk level and reward magnitude) and internal (inclination toward risk-taking) state characteristics. Throughout decision-making, presence of risk was encoded by selective signals originating in both lOFC population-level activity and responses of event-selective lOFC neuronal subpopulations. This suggests that cognitive maps during decision-making may be segregated into separate risky vs risk-free state spaces, each associated with distinct neuronal representations generated by lOFC. This information may mediate choice calculations by other brain regions more tightly linked to action selection.

During decision-making, choice represents the transition between one state and the next, and is completed when the neuronal system reaches a threshold that that initiates movement through the cognitive map. Importantly, when conflicting motivational inputs (such as reward approach or risk avoidance) are present in the environment, those inputs compete to push the current state toward transitioning to one adjacent state or another ^118–122^. Stimulus-selective attractor states often arise from strong recurrent excitation within a local cortical circuit ^121,123–125^, which mirrors the encoding of large and small reward magnitudes by distinct subpopulations’ selective excitation observed in the current work. Critically, the increased large reward-associated lOFC activation in risk-preferring rats may bias these subjects toward this option despite the presence of risk. Abnormal lOFC activity is observed in many psychiatric disorders ^36,42–44,47^; this may engender suboptimal cognitive maps that fail to adequately evaluate risk and reward, causing persistent and potentially maladaptive risky decision-making ^6,103,104,110,126^.

Machine learning models revealed that lOFC activity can be used to predict both risk level and choice on a trial-by-trial basis. Interestingly, ability to predict choice was amplified by including information about the previous trial’s outcome into the predictive model. Thus, population-level signaling in lOFC may draw upon recent actions and outcomes to update the network and proceed through the risky decision-making process. Interestingly, accuracy of choice prediction from lOFC activity was lower for trials in which rats chose their non-preferred reward (large, risky for risk-averse and small, safe for risk-preferring rats), suggesting that lOFC guidance of choice is biased toward the preferred option. Collectively, these data suggest that lOFC functions as a hub in which active and passive cognitive processes associated with risky decision-making (i.e., guiding action selection and signaling environmental risk) converge and cooccur.

## Conclusion

The combined implication of our population-level, ensemble-specific, and trial-by-trial machine learning analyses is that the lOFC differentiates between different environmental states with distinct risk levels or reward magnitude, repeatedly signaling the presence of risk throughout the decision-making process. Furthermore, these maps of task space differ based on subjective risk-preference, suggesting that a bias toward risk-taking may arise from reduced lOFC signaling of risk in conjunction with exaggerated encoding of reward magnitude.

## Acknowledgements

The authors thank Drs. Mahsa Moaddab and Michael McDannald for technical consultation. This work was supported by R15DA046797 (NWS), F31DA050458 (DBKG), and a Community of Research Scholars Grant from the University of Memphis (NWS & MY). The authors have no conflicts of interest to disclose.

## STAR ★METHODS

### SUBJECT DETAILS

Adult, male Long-Evans rats (PND>100) were obtained from Envigo Corp and kept on a 12-hour light/dark cycle beginning with lights off at 7am. Rats had ad libitum access to water and food for one week following transportation, after which they were food restricted to 85% free feeding baseline weight to increase performance motivation.

### METHOD DETAILS

#### Behavioral Apparatus

All behavior was measured using MedAssociates (FairFax, VA) modular operant conditioning chambers equipped with a shock grate floor and two retractable levers flanking a food trough connected to a pellet dispenser. Individual lights were positioned above each lever, on the opposite wall, near the chamber ceiling, and at the top of the food trough (which extended to the ceiling of the chamber to minimize jostling of implanted electrodes). Each operant chamber is housed in a sound attenuating chamber. A rotating commutator (Plexon) extended through the ceiling toward the operant chamber and a small hole in the ceiling of the operant chamber allowed a reinforced headstage cable to connect implanted electrodes to the commutator. Custom-written MedPC codes were used to control operant chamber functioning and to record subject behavior.

#### Instrumental Shaping

Animals went through instrumental shaping before progressing to the risky decision-making task^2^. The day immediately prior to shaping, rats were placed in their assigned operant chambers for five minutes and a handful of sucrose pellets were dropped in their home cages to reduce neophobia. Shaping began with teaching rats to associate the food trough with reinforcer delivery by delivering 38 pellets one at a time every 20 ± 10s, accompanied by illumination of the food trough until pellets are collected. After subjects learned to collect delivered pellets, they progressed to lever press shaping, wherein a single lever extended and subjects could earn up to 50 reinforcers in an FR-1 lever press schedule. Each lever was trained in subsequent sessions, with the order of lever location counterbalanced across subjects. After learning to respond on each lever, subjects trained to perform an extended head entry to an empty but illuminated food trough. After holding the head entry for 1 full second, the trough light extinguished and either the left or right lever was pseudorandomly extended. A response on the extended lever caused the lever to retract, delivered a single pellet reinforcer, and reilluminated the food trough. Failure to respond within 10s of cue presentation ended the trial and led to a 30s timeout period wherein all lights were extinguished. After subjects successfully completed 35 responses on each lever, they progressed to reward magnitude discrimination. During this stage, each lever was assigned either a “large” or “small” reward identity (counterbalanced across subjects). A 1-second head entry hold pseudorandomly extended each lever in isolation 4 times. A press on the large reward lever delivered 3 food pellets while a press on the small reward lever delivered 1 pellet. After 4 extensions of each lever (for a total of 8 trials) head-entries caused both levers to extend together (“choice” trial). A press on either lever delivered the associated reward magnitude and retracted both levers, followed by a 10 ± 4s intertrial interval (ITI). Subjects trained in magnitude discrimination until they demonstrated consistent preference for the large reward-associated lever, after which they progressed to the Risky Decision-making Task.

#### Risky Decision-Making Task (RDT)

During this modified version of the previously published RDT^2,4^, rats chose between large and small rewards, with the large reward accompanied by risk of footshock punishment. Each of the two blocks of trials (0 vs 50% risk) consisted of eight forced choice trials, followed by 40 free choice trials. Forced choice trials existed to inform subjects of updated risk contingencies. During each trial, the cue light above only one of the levers was illuminated along with the food trough, and a successful trial 1 second head entry into the illuminated trough caused the trough light to terminate and extended the lever below the illuminated cue light. Levers were presented 4 times each in pseudorandom order, such that the same lever was never presented more than twice in succession.

After completion of 8 forced choice trials, rats progressed to free choice trials. These trials were comparable to forced choice, except both levers were available during the choice stage, enabling measurement of preference for safe vs risky outcomes (Fig. 1A). First, the food trough light and cue lights above both levers were illuminated, then subjects initiated the trial by performing a sustained 1-second head entry to the illuminated food trough. Enforcing this period of nonmovement prior to lever extension facilitated distinct analyses of deliberation-evoked neuronal activity that was unaffected by initiation of a choice or other movement. Upon completing this hold, food trough and lever-cue lights were extinguished, and levers extended on either side of the food trough. A response on either lever retracted both levers, illuminated the food trough, and after a 0.5s delay delivered either a small (1 pellet) or large (3 pellets) reward. The large reward was accompanied with a risk of mild (0.3mA), 1-second footshock punishment delivered concurrently with either 0% (block 1) or 50% (block 2) of large reward choices. The order of risk blocks (ascending risk: 0%→ 50% or descending risk: 50% → 0%) was counterbalanced across subjects. A 10 ± 4s ITI began 15s after food delivery or upon food collection, whichever occurred first.

In both forced and free choice trials, failure to begin the initiation hold or press a lever within 15s of stimulus presentation immediately began an ITI and marked the trial as an omission. Premature withdrawal from the food trough during the 1s trial initiation hold also immediately began an ITI and was recorded as a failure. Subjects were required to complete each trial by pressing a lever in order to progress through the task. Response omission or initiation failure forced subjects to repeat the uncompleted trial after the ITI. This trial-completion criteria ensured that 1) subjects were exposed to the current risk contingencies within each block, and 2) there were enough free choice trials to limit the trial X trial variability in electrophysiological data obtained during behavior.

#### Electrode Implantation Surgery

After training in RDT, rats were implanted with drivable, 16-channel tungsten microelectrode arrays (Innovative Neurophysiology, Durham, NC) for single unit recording. Electrodes were implanted unilaterally above lOFC (+3.24 AP, +3.0 ML, −5.5. DV from skull)^128^. Immediately after surgery, the array was advanced 0.25mm from the housing cannula. After surgery, rats were restored to free-feeding schedule for five days, followed by re-establishment of food restriction and three days of habituation to the headstage cable. They then performed in RDT while attached to the headstage cable until they consistently completing trials in both risk contingencies. Electrodes were initially positioned 1.0mm above lOFC (-4.5 DV). Once consistent behavior was obtained, electrodes were advanced to the target region (-5.0 DV) and neural activity was recorded during performance in RDT. Electrodes were advanced again before each new session of RDT and neural activity was recorded between ranges of -5.0 to -6.0 DV from skull.

#### Recording During Behavior

Single-unit recording measures fluctuations in electrical activity from adjacent neurons with microelectrodes implanted into specific brain regions of an intact rat ^61,66,129^. Electrical signals were buffered by a headstage amplifier, then amplified and analog band pass filtered via preamplifier. Lightweight and unobtrusive electrode arrays, and a rotating commutator (Plexon, Dallas, TX) connected to a reinforced headstage cable ensure free movement. Critically, previous studies have found that comparable electrodes, headstage cable, and commutators do not alter action latency or task engagement in either adult or smaller adolescent rats ^66,128,130,131^. Spikes were analog filtered between 300 Hz and 8 kHz and digitized at 40 kHz. The MedPC behavioral system controller sent TTL pulses to the neural data acquisition system to synchronize behavioral events with neural data. After data collection, electrical activity was manually sorted into single-units according to standardized techniques using spike waveform topology ^66,129,132^ in Plexon Offline Sorter.

#### Histology

At the end of the experiment, rats were euthanized with chloral hydrate, then perfused with saline and 10% formalin solution. Brains were stored in formalin solution until sectioned. For at least 24 hours before sectioning, brains were soaked in a sucrose solution. They were then sectioned into 60-μm coronal slices using a MicroTome cryostat and mounted to slides. Electrode placements was verified under a light microscope ^66,128^

### QUANTIFICATION AND STATISTICAL ANALYSIS

#### Neuronal Population and Subpopulation Statistical Analysis Plan

Prior to electrode implantation, acquisition of RDT was demonstrated by consistent completion of 40 choice trials in both the 0% and 50% risk conditions. Following surgery, risky decision-making of each subject was averaged across sessions from which neuronal data was analyzed. Geometric area under the curve (AUC) was calculated using percent large reward choice in both blocks of RDT. A one-way ANOVA used this AUC as a general measure of risk preference to compare behavior between ascending and descending versions of RDT. Separate Repeated-Measures ANOVAs were used to examine the effect of risk on preference for the large reward and latency to physically choose the large reward via lever press.

Isolated single unit data was analyzed in Neuroexplorer (Plexon, Dallas, TX) and with custom written Matlab functions (MathWorks, Nattick, MA). Neuronal spikes during choice trials were binned (50ms bins), smoothed with a 5-point moving rectangular kernel, and Z-score normalized relative to their activity during a 5 second period of the ITI. This normalized activity was examined for units responsive to the 3 stages of the decision-making process (Fig 1A): deliberation, action selection, and outcome anticipation. Deliberation was measured during the 1s initiation hold, beginning at trough entry and ending at lever extension. Action selection was defined as in the quarter second period immediately preceding lever press. Outcome anticipation was defined as the 0.5s period between choice and outcome delivery, beginning at lever press and ending at reinforcer delivery.

Neural data was investigated during each event of interest using a 3-step statistical analysis plan. First, mean population activity was calculated for each stage of decision-making. A unit was defined as responding to an event if its firing rate during the event was altered (either increased or decreased) from its baseline firing rate (Table 1). Overall neuronal engagement was quantified as the percent of recorded units which were selectively suppressed or activated during a trial stage. We further qualified neuronal engagement by calculating the ratio of activated/suppressed units. Separate χ^2^ tests for independence were used to test for differences between risk conditions and choice behavior in both neuronal engagement (responsive vs non-responsive unit counts) and the activated/suppressed ratio (activated vs suppressed vs non-responsive unit counts) during each stage of decision making. A Repeated-Measures ANOVA compared net population activity between risk conditions at each stage of decision-making.

The second step of our statistical analysis plan assessed whether population-level analyses obscured neuronal encoding of RDT due to activated and suppressed neurons canceling each other out. We conducted separate group analyses on selectively responsive neuronal units. Units were categorized as activated or suppressed if they met 2 criteria: 1) a change in average z-score firing rate (either an increase or decrease from baseline) during the stage window, and 2) duration of change in firing rate sustained over consecutive 50ms bins. Exact criteria for magnitude and duration of change were customized to each stage (Table 1).

The third step of our statistical analysis plan further parsed apart neuronal encoding of RDT by comparing how modulations in firing rate/response topography differed between trials with 0% vs 50% risk, between trials in which subjects selected the large vs small reward, and how subjective risk-preference corresponds to differences in task-evoked neuronal responses. Repeated-Measures ANOVAs were used to compare risk-taking vs risk-averse rats’ neuronal firing rates within specific task stages and between different risk levels when choosing large rewards, with the goal of determining how neuronal circuitry drives a subset of rats toward persistent risk-taking or aversion.

#### Machine Learning Analyses

The large, variable dataset gathered through this technique makes it optimal for advanced computational techniques to determine predictive relationships between features of the neuronal data and behavior. Random Forests (RF) models use a collection of decision trees to classify an observation in the original dataset^60^. First, a decision-tree is grown by splitting the dataset into 2 or more sets according to the feature which provides the most homogenous segregation of datapoints into the different classes. Each of these sets are then re-split using the remaining available features. This is repeated until all features have been used, at which point the decision-tree is “fully grown”. Fully grown decision-trees are excellent at capturing complex interactions within the data but are exceptionally noisy. Random forests minimize the variance of decision-trees by averaging a collection of de-correlated trees, a process known as bagging. Each bagged tree is then included in the forest, assigned a classification based on the average of its aggregate trees, and given a vote for that class. The RF then attempts classifying observations from the original dataset based on majority vote of the forest and calculates its accuracy based on these predictions. These models are extremely useful in identifying the most important features in classifying the dataset as well as determining the predictive quality of any given feature^133^–^135^.

Here, we used RF models to investigate features of neuronal activity and determine which were most relevant for encoding risky decision-making. The models were trained and tested using firing rates of each recorded unit on a trial-by-trial basis to classify a specific trial according to 1) choice behavior and 2) risk level from ensemble activity. Thus, there were 2 distinct classes which each model tried to classify: 1) safe block and risky block, or 2) safe choice trials or risky choice trials. The performance of the machine learning model was quantified using 4 distinct classification metrics: Accuracy, Precision, Recall, and F1-score (Supplemental Table 2). Performance metrics were deemed stable if they exhibited <5% variance within trial stage. Differences in lOFC encoding based on subjective risk preference were analyzed using f1-score, which is the harmonic mean of precision and recall and provides a more normalized metric to compared between groups. Within each risk group, differences in precision and recall were further compared at the level of discrete classes for each risk group. Separate Repeated Measures ANOVAs for risk-preferring and risk-averse rats were used assessed differences in lOFC encoding (precision and recall) when classifying risky choice trials vs when classifying safe choice trials. To test whether Random Forest decoding of lOFC activity during risky decision-making is sufficient to predict behavior, a linear regression was conducted to predict the average number of risky and safe choices by each risk-group, using Precision, Recall, and F1-score for independent variables. Notably, class (whether a given metric corresponds to risky or safe choice trials) and risk group were excluded to avoid biasing the regression model. Principal Components Analysis assessed variance in each risk group’s precision, recall, and f-1 score for each class. Direct oblimin rotation was conducted to aid interpretation. Logistic regression used the Random Forest performance metrics precision and recall for each case (risky choice and safe choice trials) from each trial stage to predict risk group.

---

**Supplemental 1.**
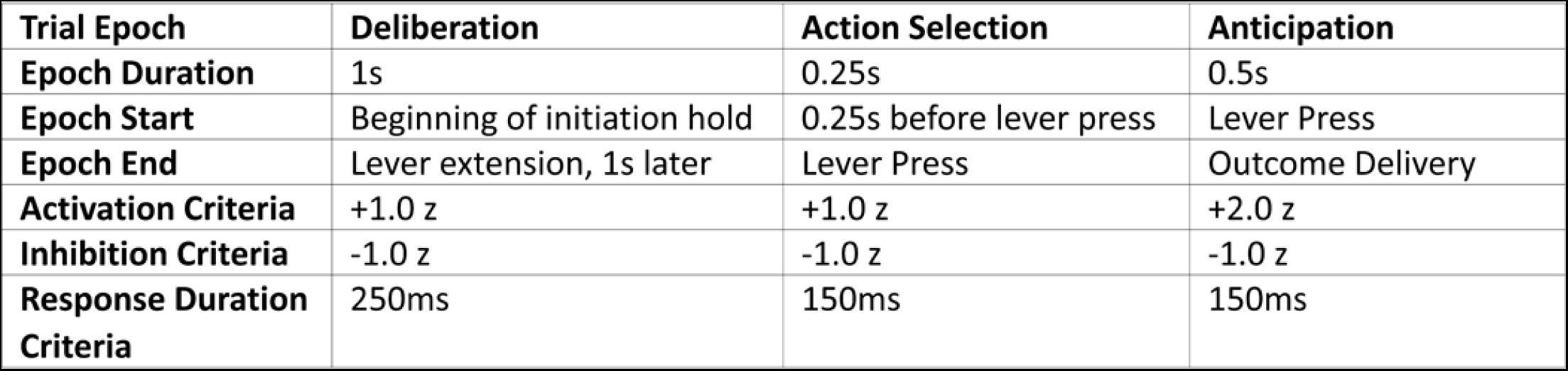
Defining events of interest during Risky Decision-making.

**Supplemental 2.**
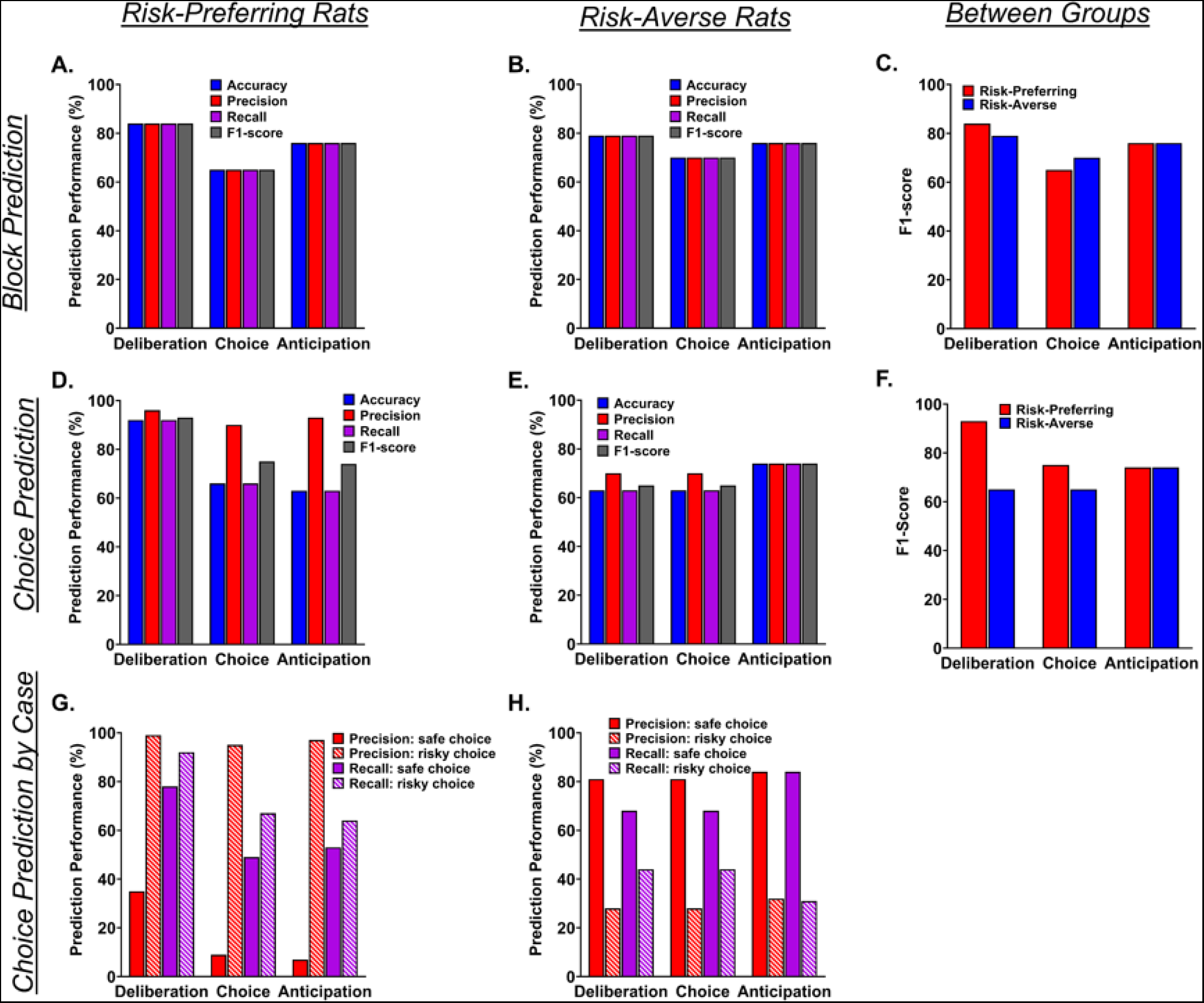
Random Forest decoding metrics within and between risk-preferring and risk-averse rats. **A-C.** lOFC activity was decoded by a parameter-optimized Random Forest (RF) model to predict whether a given trial occurred within the 0% risk or 50% risk condition of RDT. In both risk-preferring (**A.**) and risk-averse (**B.**) rats, lOFC encoded the presence/absence of risk in the environment above chance and consistently across each stage of decision-making. Subjective risk preference did not correlate with strong differences in model performance (**C.**). **D-H.** lOFC encoding of choice between large, risky vs small, safe rewards was less consistent between and within risk-groups. In risk-preferring rats (**D.**), all RF metrics except precision dropped significantly during choice and anticipation, indicating that in risk-preferring rats lOFC encoding may be more relevant in comparing upcoming choice options, rather than in directing action selection or anticipation. In risk-averse rats (**E.**) precision was elevated above other metrics during deliberation and choice, but all metrics rose to equal levels during anticipation, indicating that in risk-averse rats lOFC encoding may be more relevant in generating predictions for upcoming events. This difference in prioritization of lOFC encoding between risk groups is reflected in less consistent model performance between risk groups (**F.**), particularly during deliberation. This discrepancy between risk groups is largely influenced by differences in lOFC encoding of preferred vs non-preferred choice trials (**G,H**). RF precision and recall dropped drastically in trials where risk-preferring rats chose small, safe rewards (**G.**) and where risk-averse rats chose large, risky reward Furthermore, precision and recall varied much more drastically for risk-preferring rats (**G.**) than for risk-averse rats (**H.**), indicating a that excessive risk preference may be related to an inefficiency in lOFC encoding during risky decision-making.

**Supplemental 3.**
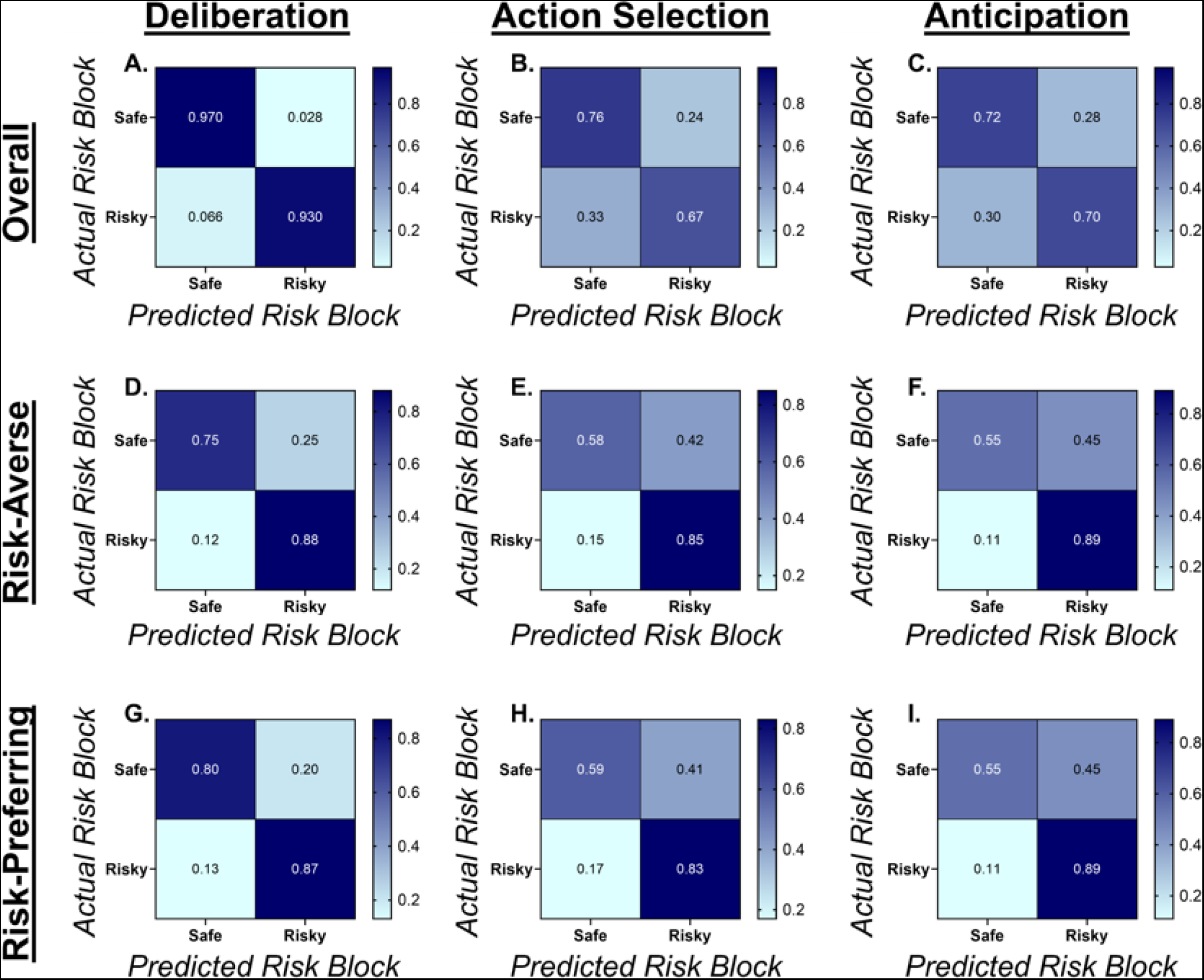
Populations activity decoding to predict current risk conditions on a trial-by-trial basis. Confusion matrices for RF classification for each trial as belonging in the 0% or 50% risky block of RDT in deliberation (col 1), action selection (col 2) and anticipation (col 3). The generalized model misclassified very few trials during deliberation (**A.**). Confusion matrices also did not strongly differ between risk-averse (**D.**) or risk-preferring (**G.**) rats during deliberation. More trials were misclassified during action selection (**B.**) and anticipation (**C.**), with a slight bias toward classifying trials as occurring in the safe block during action selection. Increased misclassifications occurred during action selection and anticipation for both risk-averse (**E,F**) and risk-preferring (**H,I**) rats. Interestingly, when parsing RF classifications by subjective risk preferences, lOFC activity appears biased toward encoding trials as occurring in the risky block, and this bias was present during both action selection and anticipation.

**Supplemental 4.**
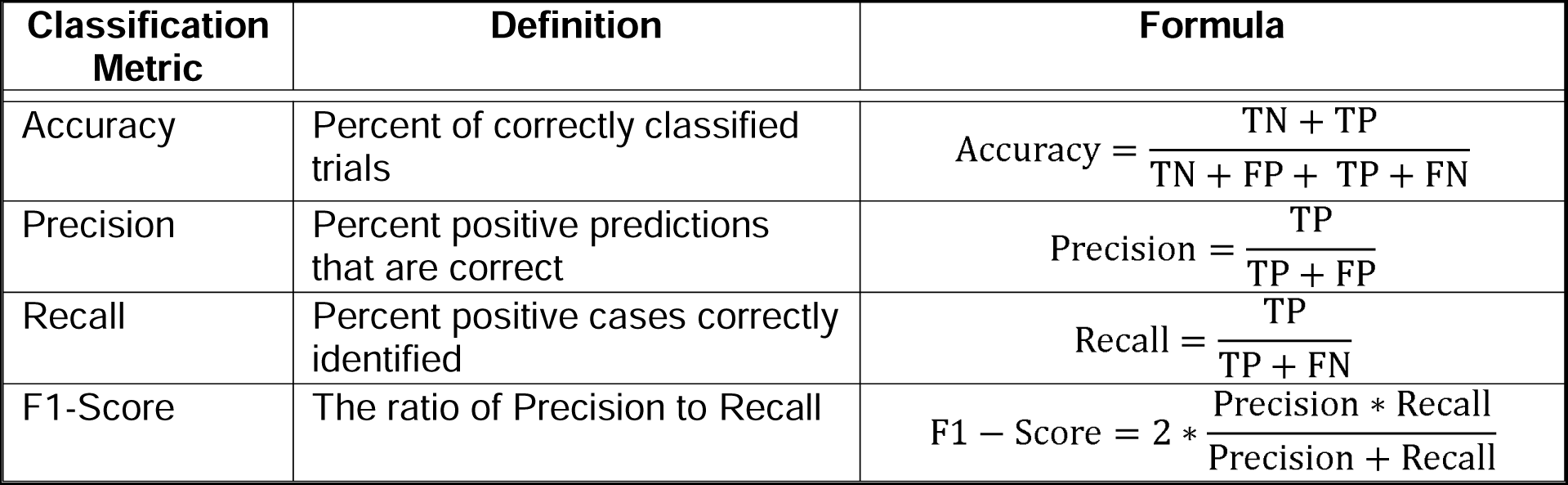
Machine Learning Classification Metrics. TN = True negative predictions. TN = True positive predictions. FN = False negative predictions. FP = False positive predictions.

## References

1. Zhao, Q., Li, H., Hu, B., Wu, H., and Liu, Q. (2017). Abstinent heroin addicts tend to take risks: ERP and source localization. Front. Neurosci. 11, 1–12. 10.3389/fnins.2017.00681.

2. Orsini, C.A., and Simon, N.W. (2020). Reward/Punishment-Based Decision Making in Rodents. Curr. Protoc. Neurosci. 93, 1–21. 10.1002/cpns.100.

3. Simon, N.W., Gilbert, R.J., Mayse, J.D., Bizon, J.L., and Setlow, B. (2009). Balancing Risk and Reward: A Rat Model of Risky Decision Making. Neuropsychopharmacology 34, 2208–2217. 10.1038/npp.2009.48.

4. Simon, N.W., and Setlow, B. (2012). Modeling Risky Decision Making in Rodents. Methods Mol Biol. 829, 165–175. 10.1007/978-1-61779-458-2.

5. Guttman, Z., Moeller, S.J., and London, E.D. (2018). Neural Underpinnings of Maladaptive Decision-Making in Addictions. Pharmacol Biochem Behav, 84–98. 10.1016/j.pbb.2017.06.014.Neural.

6. Verdejo-García, A., Chong, T.T.-J., Stout, J.C., Yucel, M., London, E.D., Yücel, M., and London, E.D. (2018). Stages of dysfunctional decision-making in addiction. Pharmacol. Biochem. Behav. 164, 99–105. 10.1016/j.pbb.2017.02.003.

7. Ferland, J.N., and Winstanley, C.A. (2016). Risk-preferring rats make worse decisions and show increased incubation of craving after cocaine. Addict. Biol. 122, 991–1001. 10.1111/adb.12388.

8. Lane, S.D., and Cherek, D.R. (2000). Analysis of risk taking in adults with a history of high risk behavior. Drug Alcohol Depend. 60, 179–187. 10.1016/S0376-8716(99)00155-6.

9. Brand, M., Roth-Bauer, M., Driessen, M., and Markowitsch, H.J. (2008). Executive functions and risky decision-making in patients with opiate dependence. Drug Alcohol Depend. 97, 64–72. 10.1016/j.drugalcdep.2008.03.017.

10. Bechara, A., Dolan, S., Denburg, N., Hindes, A., Anderson, S.W., and Nathan, P.E. (2001). Decision-making deficits, linked to a dysfunctional ventromedial prefrontal cortex, revealed in alcohol and stimulant abusers. Neuropsychologia 39, 376–389. 10.1016/S0028-3932(00)00136-6.

11. Brand, M., Labudda, K., and Kalbe, E. (2004). Decision-making impairments in patients with Parkinson’s disease. 15, 77–85.

12. Toplak, M.E., Sorge, G.B., Benoit, A., West, R.F., and Stanovich, K.E. (2010). Decision-making and cognitive abilities: A review of associations between Iowa Gambling Task performance, executive functions, and intelligence. Clin. Psychol. Rev. 30, 562–581. 10.1016/j.cpr.2010.04.002.

13. Toplak, M.E., Liu, E., Macpherson, R., Toneatto, T., and Stanovich, K.E. (2007). The reasoning skills and thinking dispositions of problem gamblers: A dual-process taxonomy. J. Behav. Decis. Mak. 20, 103–124. 10.1002/bdm.544.

14. Winstanley, C.A., Eagle, D.M., and Robbins, T.W. (2006). Behavioral models of impulsivity in relation to ADHD: Translation between clinical and preclinical studies. Clin. Psychol. Rev. 26, 379–395. 10.1016/j.cpr.2006.01.001.

15. Koffarnus, M.N., and Kaplan, B.A. (2018). Clinical models of decision making in addiction. Pharmacol. Biochem. Behav. 164, 71–83. 10.1016/j.pbb.2017.08.010.

16. Lauriola, M., Panno, A., Levin, I.P., and Lejuez, C.W. (2014). Individual Differences in Risky Decision Making: A Meta-analysis of Sensation Seeking and Impulsivity with the Balloon Analogue Risk Task. J. Behav. Decis. Mak. 27, 20–36. 10.1002/bdm.1784.

17. Gabriel, D.B.K., Freels, T.G., Setlow, B., and Simon, N.W. (2019). Risky decision-making is associated with impulsive action and sensitivity to first-time nicotine exposure. Behav. Brain Res. 10.1016/j.bbr.2018.10.008.

18. Liley, A.E., Joyner, H., Gabriel, D.B.K., and Simon, N.W. (2022). Effects of the Psychoactive Compounds in Green Tea on Risky Decision-making. Behav. Pharmacol. 33, 32–41.

19. Shimp, K.G., Mitchell, M.R., Beas, B.S., Bizon, J.L., and Setlow, B. (2015). Affective and cognitive mechanisms of risky decision making. Neurobiol. Learn. Mem. 117, 60–70. 10.1016/j.nlm.2014.03.002.

20. Orsini, C.A., Willis, M.L., Gilbert, R.J., Bizon, J.L., and Setlow, B. (2016). Sex differences in a rat model of risky decision making. Behav. Neurosci. 130, 50–61. 10.1037/bne0000111.

21. Mitchell, M.R., Weiss, V.G., Beas, B.S., Morgan, D., Bizon, J.L., and Setlow, B. (2014). Adolescent risk taking, cocaine self-administration, and striatal dopamine signaling. Neuropsychopharmacology 39, 955–962. 10.1038/npp.2013.295.

22. Simon, N.W., Montgomery, K.S., Beas, B.S., Mitchell, M.R., LaSarge, C.L., Mendez, I.A., Banuelos, C., Vokes, C.M., Taylor, A.B., Haberman, R.P., et al. (2011). Dopaminergic modulation of risky decision-making. J. Neurosci. 31, 17460–17470. 10.1523/JNEUROSCI.3772-11.2011.

23. Orsini, C.A., Blaes, S.L., Dragone, R.J., Betzhold, S.M., Finner, A.M., Bizon, J.L., and Setlow, B. (2020). Distinct relationships between risky decision making and cocaine self-administration under short-and long-access conditions. Prog. Neuro-Psychopharmacology Biol. Psychiatry 98, 109791. 10.1016/j.pnpbp.2019.109791.

24. Olshavsky, M.E., Shumake, J., Rosenthal, A.A., Kaddour-Djebbar, A., Gonzalez-Lima, F., Setlow, B., and Lee, H.J. (2014). Impulsivity, risk-taking, and distractibility in rats exhibiting robust conditioned orienting behaviors. J. Exp. Anal. Behav. 102, 162–178. 10.1002/jeab.104.

25. Gabriel, D.B.K., Liley, A.E., Franks, H.T., Minnes, G.L., Tutaj, M., Dwinell, M.R., de Jong, T. V., Williams, R.W., Mulligan, M.K., Chen, H., et al. (2023). Divergent risky decision-making and impulsivity behaviors in Lewis rat substrains with low genetic difference. Behav. Neurosci. 10.1037/BNE0000557.

26. Freels, T.G., Gabriel, D.B.K., Lester, D.B., and Simon, N.W. (2019). Risky Decision-Making Predicts Dopamine Release Dynamics in Nucleus Accumbens Shell. bioRxiv, 572263. 10.1101/572263.

27. Bronner, R. (2003). Pathologies of Decision-makingl:Causes, Forms, and Handling. MIR Manag. Int. Rev. 43, 85–101.

28. Montague, P.R., and Berns, G.S. (2002). Neural economics and the biological substrates of valuation. Neuron 36, 265–284. 10.1016/S0896-6273(02)00974-1.

29. Dohmen, T., Falk, A., Huffman, D., Sunde, U., Schupp, J., and Wagner, G.G. (2011). Individual risk attitudes: Measurement, determinants, and behavioral consequences. J. Eur. Econ. Assoc. 9, 522–550. 10.1111/j.1542-4774.2011.01015.x.

30. Breysse, E., Meffre, J., Pelloux, Y., Winstanley, C.A., and Baunez, C. (2021). Decreased risk-taking and loss-chasing after subthalamic nucleus lesion in rats. Eur. J. Neurosci. 53, 2362–2375. 10.1111/ejn.14895.

31. Stalnaker, T.A., Raheja, N., and Schoenbaum, G. (2021). Orbitofrontal State Representations Are Related to Choice Adaptations and Reward Predictions. Behavioral/Cognitive 41, 1941–1951.

32. Wilson, R.C., Takahashi, Y.K., Schoenbaum, G., and Niv, Y. (2014). Orbitofrontal cortex as a cognitive map of task space. Neuron 81, 267–279. 10.1016/j.neuron.2013.11.005.Orbitofrontal.

33. Hosokawa, T., Kato, K., Inoue, M., and Mikami, A. (2007). Neurons in the macaque orbitofrontal cortex code relative preference of both rewarding and aversive outcomes. Neurosci. Res. 57, 434–445. 10.1016/j.neures.2006.12.003.

34. Schoenbaum, G., and Roesch, M. (2005). Orbitofrontal cortex, associative learning, and expectancies, 10.1016/j.neuron.2005.07.018.

35. Plassmann, H., O’Doherty, J., and Rangel, A. (2007). Orbitofrontal cortex encodes willingness to pay in everyday economic transactions. J. Neurosci. 27, 9984–9988. 10.1523/JNEUROSCI.2131-07.2007.

36. Kohno, M., Morales, A.M., Ghahremani, D.G., Hellemann, G., and London, E.D. (2014). Risky decision making, prefrontal cortex, and mesocorticolimbic functional connectivity in methamphetamine dependence. JAMA Psychiatry 71, 812–820. 10.1001/jamapsychiatry.2014.399.

37. Orsini, C.A., Trotta, R.T., Bizon, J.L., and Setlow, B. (2015). Dissociable Roles for the Basolateral Amygdala and Orbitofrontal Cortex in Decision-Making under Risk of Punishment. J. Neurosci. 35, 1368–1379. 10.1523/JNEUROSCI.3586-14.2015.

38. Pais-Vieira, M., Lima, D., and Galhardo, V. (2007). Orbitofrontal cortex lesions disrupt risk assessment in a novel serial decision-making task for rats. Neuroscience 145, 225–231. 10.1016/j.neuroscience.2006.11.058.

39. Zeeb, F.D., Floresco, S.B., and Winstanley, C.A. (2010). Contributions of the orbitofrontal cortex to impulsive choice: interactions with basal levels of impulsivity, dopamine signalling, and reward-related cues. Psychopharmacology (Berl). 211, 87–98. 10.1007/S00213-010-1871-2.

40. Fuchs, R.A., Evans, K.A., Parker, M.P., and See, R.E. (2004). Behavioral/Systems/Cognitive Differential Involvement of Orbitofrontal Cortex Subregions in Conditioned Cue-Induced and Cocaine-Primed Reinstatement of Cocaine Seeking in Rats. 10.1523/JNEUROSCI.1924-04.2004.

41. St. Onge, J.R., and Floresco, S.B. (2010). Prefrontal cortical contribution to risk-based decision making. Cereb. Cortex 20, 1816–1828. 10.1093/cercor/bhp250.

42. Bolla, K.I., Eldreth, D.A., London, E.D., Kiehl, K.A., Mouratidis, M., Contoreggi, C., Matochik, J.A., Kurian, V., Cadet, J.L., Kimes, A.S., et al. (2003). Orbitofrontal cortex dysfunction in abstinent cocaine abusers performing a decision-making task. Neuroimage 19, 1085–1094. 10.1016/S1053-8119(03)00113-7.

43. Pascoli, V., Hiver, A., Van Zessen, R., Loureiro, M., Achargui, R., Harada, M., Flakowski, J., and Lüscher, C. (2018). Stochastic synaptic plasticity underlying compulsion in a model of addiction. Nature. 10.1038/s41586-018-0789-4.

44. Schoenbaum, G., Chang, C.Y., Lucantonio, F., and Takahashi, Y.K. (2016). Thinking Outside the Box: Orbitofrontal Cortex, Imagination, and How We Can Treat Addiction, 10.1038/npp.2016.147.

45. Barrus, M.M., Hosking, J.G., Cocker, P.J., and Winstanley, C.A. (2017). Inactivation of the orbitofrontal cortex reduces irrational choice on a rodent Betting Task. Neuroscience 345, 38–48. 10.1016/j.neuroscience.2016.02.028.

46. Duan, L.Y., Horst, N.K., Cranmore, S.A.W., Horiguchi, N., Cardinal, R.N., Roberts, A.C., and Robbins, T.W. (2021). Controlling one’s world: Identification of sub-regions of primate PFC underlying goal-directed behavior. Neuron 109, 2485–2498.e5. 10.1016/j.neuron.2021.06.003.

47. Meyer, H.C., and Bucci, D.J. (2016). Imbalanced Activity in the Orbitofrontal Cortex and Nucleus Accumbens Impairs Behavioral Inhibition. Curr. Biol. 26, 2834–2839. 10.1016/j.cub.2016.08.034.

48. Turner, K.M., Balleine, B.W., and Bradfield, L.A. (2020). Does disrupting the orbitofrontal cortex alter sensitivity to punishment? A potential mechanism of compulsivity. Behav. Neurosci. 135, 174–181. 10.1037/bne0000443.

49. Moorman, D.E. (2018). The role of the orbitofrontal cortex in alcohol use, abuse, and dependence, 10.1016/j.pnpbp.2018.01.010.

50. Padoa-Schioppa, C. (2007). Orbitofrontal cortex and the computation of economic value. Ann. N. Y. Acad. Sci. 1121, 232–253. 10.1196/annals.1401.011.

51. Orsini, C.A., Heshmati, S.C., Garman, T.S., Wall, S.C., Bizon, J.L., and Setlow, B. (2018). Contributions of medial prefrontal cortex to decision making involving risk of punishment. Neuropharmacology. 10.1016/j.neuropharm.2018.07.018.

52. Orsini, C.A., Hernandez, C.M., Bizon, J.L., and Setlow, B. (2018). Deconstructing value-based decision making via temporally selective manipulation of neural activity: Insights from rodent models. Cogn. Affect. Behav. Neurosci. 19, 459–476. 10.3758/s13415-018-00649-0.

53. Orsini, C.A., Hernandez, C.M., Singhal, S., Kelly, K.B., Frazier, C.J., Bizon, J.L., and Setlow, B. (2017). Optogenetic inhibition reveals distinct roles for basolateral amygdala activity at discrete timepoints during risky decision making. J. Neurosci. 041493, 2344–17. 10.1523/JNEUROSCI.2344-17.2017.

54. Witkowski, P.P., Park, S.A., and Boorman, E.D. (2022). Neural mechanisms of credit assignment for inferred relationships in a structured world. Neuron 110, 2680–2690.e9. 10.1016/j.neuron.2022.05.021.

55. Behrens, T.E.J., Muller, T.H., Whittington, J.C.R., Mark, S., Baram, A.B., Stachenfeld, K.L., and Kurth-Nelson, Z. (2018). What Is a Cognitive Map? Organizing Knowledge for Flexible Behavior. Neuron 100, 490–509. 10.1016/J.NEURON.2018.10.002.

56. Bradfield, L.A., and Hart, G. (2020). Rodent medial and lateral orbitofrontal cortices represent unique components of cognitive maps of task space. Neurosci. Biobehav. Rev. 108, 287–294. 10.1016/j.neubiorev.2019.11.009.

57. Tolman, E.C. (1948). Cognitive maps in rats and men. Psychol. Rev. 55, 189–208. 10.1037/H0061626.

58. Namboodiri, V.M.K., and Stuber, G.D. (2021). The learning of prospective and retrospective cognitive maps within neural circuits. Neuron 109, 3552–3575. 10.1016/j.neuron.2021.09.034.

59. Park, S.A., Miller, D.S., Nili, H., Ranganath, C., and Boorman, E.D. (2020). Map Making: Constructing, Combining, and Inferring on Abstract Cognitive Maps. Neuron 107, 1226–1238.e8. 10.1016/j.neuron.2020.06.030.

60. Leo Breiman (2001). Random Forests 10.1023/A:1010933404324.

61. Simon, N.W., and Moghaddam, B. (2015). Neural processing of reward in adolescent rodents. Dev. Cogn. Neurosci. 11, 145–154. 10.1016/j.dcn.2014.11.001.

62. Groman, S.M., Daeyeol Lee, J., and Taylor, J.R. (2021). Unlocking the Reinforcement-Learning Circuits of the Orbitofrontal Cortex. Behav Neurosci. 135, 120–128. 10.1037/bne0000414.Unlocking.

63. Sadacca, B.F., Wied, H.M., Lopatina, N., Saini, G.K., Nemirovsky, D., and Schoenbaum, G. (2018). Orbitofrontal neurons signal sensory associations underlying model-based inference in a sensory preconditioning task. Elife 7, e30373. 10.7554/eLife.30373.

64. Lucantonio, F., Gardner, M.P.H., Mirenzi, A., Newman, L.E., Takahashi, Y.K., and Schoenbaum, G. (2015). Neural estimates of imagined outcomes in basolateral amygdala depend on orbitofrontal cortex. J. Neurosci. 35, 16521–16530. 10.1523/JNEUROSCI.3126-15.2015.

65. Costa, K.M., Scholz, R., Lloyd, K., Moreno-Castilla, P., Gardner, M.P.H., Dayan, P., and Schoenbaum, G. (2023). The role of the lateral orbitofrontal cortex in creating cognitive maps. Nat. Neurosci. 26, 107–115. 10.1038/s41593-022-01216-0.

66. Simon, N.W., Wood, J., and Moghaddam, B. (2015). Action-outcome relationships are represented differently by medial prefrontal and orbitofrontal cortex neurons during action execution. J. Neurophysiol. 114, 3374–3385. 10.1152/jn.00884.2015.

67. O’Doherty, J., Kringelbach, M.L., Rolls, E.T., Hornak, J., and Andrews, C. (2001). Abstract reward and punishment representations in the human orbitofrontal cortex. Nat. Neurosci. 10.1038/82959.

68. Van Duuren, E., Lankelma, J., and Pennartz, C.M.A. (2008). Population coding of reward magnitude in the orbitofrontal cortex of the rat. J. Neurosci. 28, 8590–8603. 10.1523/JNEUROSCI.5549-07.2008.

69. Nedios, S., Konstantinos Iliodromitis, Kowalewski, C., Bollmann, A., Hindricks, · Gerhard, Dagres, N., and Harilaos Bogossian, · (2022). Big Data in electrophysiology. Herzschrittmachertherapie + Elektrophysiologie 33, 26–33. 10.1007/s00399-022-00837-z.

70. Thibeault, K.C., Kutlu, M.G., Sanders, C., and Calipari, E.S. (2019). Cell-type and projection-specific dopaminergic encoding of aversive stimuli in addiction. Brain Res. 1713, 1–15. 10.1016/j.brainres.2018.12.024.

71. Orsini, C.A., Moorman, D.E., Young, J.W., Setlow, B., and Floresco, S.B. (2015). Neural mechanisms regulating different forms of risk-related decision-making: Insights from animal models. Neurosci. Biobehav. Rev. 58, 147–167. 10.1016/j.neubiorev.2015.04.009.

72. Anderson, A.K., Christoff, K., Stappen, I., Panitz, D., Ghahremani, D.G., Glover, G., Gabrieli, J.D.E., and Sobel, N. (2003). Dissociated neural representations of intensity and valence in human olfaction. Nat. Neurosci. 6, 196–202. 10.1038/nn1001.

73. Small, D.M., Gregory, M.D., Mak, Y.E., Gitelman, D., Mesulam, M.M., and Parrish, T. (2003). Dissociation of neural representation of intensity and affective valuation in human gustation. Neuron 39, 701–711. 10.1016/S0896-6273(03)00467-7.

74. Kobayashi, S., Nomoto, K., Watanabe, M., and Hikosaka, O. (2006). Influences of Rewarding and Aversive Outcomes on Activity in Macaque Lateral Prefrontal Cortex. 861–870. 10.1016/j.neuron.2006.08.031.

75. Morrison, S.E., and Salzman, C.D. (2009). The Convergence of Information about Rewarding and Aversive Stimuli in Single Neurons. J. Neurosci. 29, 11471–11483. 10.1523/JNEUROSCI.1815-09.2009.

76. Hirokawa, J., Vaughan, A., Masset, P., Ott, T., and Kepecs, A. (2019). Frontal cortex neuron types categorically encode single decision variables. Nature 576, 446–451. 10.1038/s41586-019-1816-9.

77. Roesch, M.R., and Olson, C.R. (2004). Neuronal Activity Related to Reward Value and Motivation in Primate Frontal Cortex. Science (80-.). 304, 307–310. 10.1126/science.1093223.

78. Roesch, M.R., Calu, D.J., Burke, K.A., and Schoenbaum, G. (2007). Should I stay or should I go? Transformation of time-discounted rewards in orbitofrontal cortex and associated brain circuits. In Annals of the New York Academy of Sciences, pp. 21–34. 10.1196/annals.1390.001.

79. Tremblay, L., and Schultz, W. (2000). Modifications of Reward Expectation-Related Neuronal Activity During Learning in Primate Orbitofrontal Cortex. J. Neurophysiol. 83, 1777–2470.

80. Rolls, E.T. (2000). The orbitofrontal cortex and reward. Cereb. Cortex 10, 284–294. 10.1093/cercor/10.3.284.

81. Schoenbaum, G., Chiba, A.A., and Gallagher, M. (1998). Orbitofrontal cortex and basolateral amygdala encode expected outcomes during learning. Nat. Neurosci. 1, 155–159. 10.1038/407.

82. Howard, J.D., and Kahnt, T. (2021). To be specific: The role of orbitofrontal cortex in signaling reward identity. Behav. Neurosci. 135, 210–217. 10.1037/BNE0000455.

83. Kim, Y., Wood, J., and Moghaddam, B. (2012). Coordinated activity of ventral tegmental neurons adapts to appetitive and aversive learning. PLoS One 7. 10.1371/journal.pone.0029766.

84. Pascoli, V., Terrier, J., Hiver, A., and Lüscher, C. (2015). Sufficiency of Mesolimbic Dopamine Neuron Stimulation for the Progression to Addiction. Neuron 88, 1054–1066. 10.1016/J.NEURON.2015.10.017.

85. Ishikawa, J., Sakurai, Y., Ishikawa, A., and Mitsushima, D. (2020). Contribution of the prefrontal cortex and basolateral amygdala to behavioral decision-making under reward/punishment conflict. Psychopharmacology (Berl). 237, 639–654. 10.1007/S00213-019-05398-7/FIGURES/6.

86. Jean-Richard-Dit-Bressel, P., and McNally, G.P. (2016). Lateral, not medial, prefrontal cortex contributes to punishment and aversive instrumental learning. Learn. Mem. 23, 607–617. 10.1101/lm.042820.116.

87. Sutton, R.S., and Barto, A.G. (2018). Reinforcement learning: An Introduction 2nd ed. (MIT press).

88. Yoo, S.B.M., and Hayden, B.Y. (2020). The Transition from Evaluation to Selection Involves Neural Subspace Reorganization in Core Reward Regions. Neuron 105, 712–724.e4. 10.1016/j.neuron.2019.11.013.

89. Zhou, J., Zong, W., Jia, C., Gardner, M.P.H., and Schoenbaum, G. (2021). Prospective representations in rat orbitofrontal ensembles. Behav. Neurosci. 10.1037/bne0000451.

90. Stalnaker, T.A., Cooch, N.K., McDannald, M.A., Liu, T.L., Wied, H., and Schoenbaum, G. (2014). Orbitofrontal neurons infer the value and identity of predicted outcomes. Nat. Commun. 5, 1–13. 10.1038/ncomms4926.

91. Berns, G.S., Chappelow, J., Cekic, M., Zink, C.F., Pagnoni, G., and Martin-Skurski, M.E. (2006). Neurobiological Substrates of Dread. Science 312, 754. 10.1126/SCIENCE.1123721.

92. Roesch, M.R., Taylor, A.R., and Schoenbaum, G. (2006). Encoding of Time-Discounted Rewards in Orbitofrontal Cortex Is Independent of Value Representation. Neuron. 10.1016/j.neuron.2006.06.027.

93. Somogyi, P., Bolam, J.P., and Smith, A.D. (1981). Monosynaptic cortical input and local axon collaterals of identified striatonigral neurons. A light and electron microscopic study using the Golgi-peroxidase transport-degeneration procedure. J. Comp. Neurol. 195, 567–584. 10.1002/CNE.901950403.

94. Haber, S.N. (2016). Corticostriatal circuitry. Dialogues Clin. Neurosci. April.

95. Floresco, S.B., St. Onge, J.R., Ghods-Sharifi, S., and Winstanley, C.A. (2008). Cortico-limbic-striatal circuits subserving different forms of cost-benefit decision making. Cogn. Affect. Behav. Neurosci. 8, 375–389. 10.3758/CABN.8.4.375.

96. Cardinal, R.N. (2006). Neural systems implicated in delayed and probabilistic reinforcement. Neural Networks 19, 1277–1301. 10.1016/j.neunet.2006.03.004.

97. Burton, A.C., Kashtelyan, V., Bryden, D.W., and Roesch, M.R. (2014). Increased firing to cues that predict low-value reward in the medial orbitofrontal cortex. Cereb. Cortex 24, 3310–3321. 10.1093/cercor/bht189.

98. Roesch, M.R., Bryden, D.W., Cerri, D.H., Haney, Z.R., and Schoenbaum, G. (2012). Willingness to wait and altered encoding of time-discounted reward in the orbitofrontal cortex with normal aging. J. Neurosci. 32, 5525–5533. 10.1523/JNEUROSCI.0586-12.2012.

99. Takahashi, Y.K., Roesch, M.R., Stalnaker, T.A., Haney, R.Z., Calu, J., Taylor, A.R., Burke, K.A., and Schoenbaum, G. (2009). The orbitofrontal cortex and ventral tegmental area are necessary for Learning From Unexpected Outcomes. Neuron 62, 269–280. 10.1016/j.neuron.2009.03.005.The.

100. Mitchell, M.R., Vokes, C.M., Blankenship, A.L., Simon, N.W., and Setlow, B. (2011). Effects of acute administration of nicotine, amphetamine, diazepam, morphine, and ethanol on risky decision-making in rats. Psychopharmacology (Berl). 218, 703–712. 10.1007/s00213-011-2363-8.

101. Roitman, J.D., and Roitman, M.F. (2010). Risk-preference differentiates orbitofrontal cortex responses to freely chosen reward outcomes. Eur. J. 2Neuroscience 31, 1492–1500. 10.1111/j.1460-9568.2010.07169.x.

102. Moorman, D.E., and Aston-jones, G. (2014). Orbitofrontal Cortical Neurons Encode Expectation-Driven Initiation of Reward-Seeking. J. Neurosci. 34, 10234–10246. 10.1523/JNEUROSCI.3216-13.2014.

103. Calaminus, C., and Hauber, W. (2008). Guidance of instrumental behavior under reversal conditions requires dopamine D1 and D2 receptor activation in the orbitofrontal cortex. Neuroscience. 10.1016/j.neuroscience.2008.04.046.

104. Rich, E.L., Stoll, F.M., and Rudebeck, P.H. (2018). Linking dynamic patterns of neural activity in orbitofrontal cortex with decision making. Curr. Opin. Neurobiol. 49, 24–32. 10.1016/j.conb.2017.11.002.

105. Rich, E.L., and Wallis, J.D. (2016). Decoding subjective decisions from orbitofrontal cortex. Nat. Neurosci. 19, 973–980. 10.1038/nn.4320.

106. Balleine, B.W., and O’Doherty, J.P. (2010). Human and rodent homologies in action control: Corticostriatal determinants of goal-directed and habitual action, 10.1038/npp.2009.131.

107. Gremel, C.M., and Costa, R.M. (2013). Orbitofrontal and striatal circuits dynamically encode the shift between goal-directed and habitual actions. Nat. Commun. 4, 1–12. 10.1038/ncomms3264.

108. Hernandez, J.S., Binette, A.N., Rahman, T., Tarantino, J.D., and Moorman, D.E. (2020). Chemogenetic Inactivation of Orbitofrontal Cortex Decreases Cue-Induced Reinstatement of Ethanol and Sucrose Seeking in Male and Female Wistar Rats. Alcohol Clin Exp Res 44, 1769–1782. 10.1111/acer.14407.Chemogenetic.

109. Arinze, I., and Moorman, D.E. (2020). Selective impact of lateral orbitofrontal cortex inactivation on reinstatement of alcohol seeking in male Long-Evans rats. Neuropharmacology 168. 10.1016/j.neuropharm.2020.108007.

110. Groman, S.M., Keistler, C., Keip, A.J., Hammarlund, E., DiLeone, R.J., Pittenger, C., Lee, D., and Taylor, J.R. (2019). Orbitofrontal Circuits Control Multiple Reinforcement-Learning Processes. Neuron, 1–13. 10.1016/j.neuron.2019.05.042.

111. Rich, E.L., and Shapiro, M. (2009). Rat prefrontal cortical neurons selectively code strategy switches. J. Neurosci. 29, 7208–7219. 10.1523/JNEUROSCI.6068-08.2009.

112. Wikenheiser, A.M., and Schoenbaum, G. (2016). Over the river, through the woods: Cognitive maps in the hippocampus and orbitofrontal cortex. Nat. Rev. Neurosci. 17, 513–523. 10.1038/NRN.2016.56.

113. Gardner, M.P.H., and Schoenbaum, G. (2021). The orbitofrontal cartographer. Behav. Neurosci. 135, 267–276. 10.1037/BNE0000463.

114. Gardner, M.P.H., Conroy, J.S., Shaham, M.H., Styer, C. V., and Schoenbaum, G. (2017). Lateral Orbitofrontal Inactivation Dissociates Devaluation-Sensitive Behavior and Economic Choice. Neuron 96, 1192–1203.e4. 10.1016/j.neuron.2017.10.026.

115. Miller, K.J., Botvinick, M.M., and Brody, C.D. (2022). Value representations in the rodent orbitofrontal cortex drive learning, not choice. Elife 11, 1–27. 10.7554/eLife.64575.

116. Gardner, M.P.H., Sanchez, D., Conroy, J.C., Wikenheiser, A.M., Zhou, J., and Schoenbaum, G. (2020). Processing in Lateral Orbitofrontal Cortex Is Required to Estimate Subjective Preference during Initial, but Not Established, Economic Choice. Neuron 108, 526–537.e4. 10.1016/j.neuron.2020.08.010.

117. Schuck, N.W., Cai, M.B., Wilson, R.C., and Niv, Y. (2016). Human Orbitofrontal Cortex Represents a Cognitive Map of State Space. Neuron 91, 1402–1412. 10.1016/j.neuron.2016.08.019.Human.

118. Enel, P., Wallis, J.D., and Rich, E.L. (2020). Stable and dynamic representations of value in the prefrontal cortex. Natl. Institutes Heal. Elife, 9:e54313. 10.7554/eLife.54313.

119. Khona, M., and Fiete, I.R. (2022). Attractor and integrator networks in the brain. Nat. Rev. Neurosci. 23, 744–766. 10.1038/s41583-022-00642-0.

120. Kriener, B., Chaudhuri, R., and Fiete, I.R. (2020). Robust parallel decision-making in neural circuits with nonlinear inhibition. PNAS 117, 25505–25516. 10.1073/pnas.1917551117.

121. Xiaojing, W. (2008). Decision Making in Recurrent Neuronal Circuits. Neuron 60, 215–234. 10.1016/j.neuron.2008.09.034.Decision.

122. Roitman, J.D., and Shadlen, M.N. (2002). Response of neurons in the lateral intraparietal area during a combined visual discrimination reaction time task. J. Neurosci. 22, 9475–9489. 10.1523/jneurosci.22-21-09475.2002.

123. Amit, D.J. (1995). The Hebbian paradigm reintegrated: Local reverberations as internal representations. Behav. Brain Sci. 18, 617–657. 10.1017/S0140525X0004022X.

124. Goldman-Rakic, P.S. (1995). Cellular basis of working memory. Neuron 14, 477–485. 10.1016/0896-6273(95)90304-6.

125. Wang, X.-J. (2001). Synaptic reverberation underlying mnemonic persistent activity. TRENDS Neurosci. 24, 455–463.

126. Rudebeck, P.H., and Rich, E.L. (2018). Orbitofrontal cortex. Curr. Biol. 28, R1083–R1088. 10.1016/j.cub.2018.07.018.

128. Paxinos, G., and Watson, C. (1997). The Rat Brain in Stereotaxic Coordinates. Acad. Press. San Diego 3rd.

129. Wood, J., Simon, N.W., Koerner, F.S., Kass, R.E., and Moghaddam, B. (2017). Networks of VTA Neurons Encode Real-Time Information about Uncertain Numbers of Actions Executed to Earn a Reward. Front. Behav. Neurosci. 11. 10.3389/fnbeh.2017.00140.

130. Kim, Y., Simon, N.W., Wood, J., and Moghaddam, B. (2016). Reward anticipation is encoded differently by adolescent ventral tegmental area neurons. Biol. Psychiatry 79, 878–886. 10.1038/s41598-019-39414-9.

131. Totah, N.K.B., Kim, Y.B., Homayoun, H., and Moghaddam, B. (2009). Anterior Cingulate Neurons Represent Errors and Preparatory Attention within the Same Behavioral Sequence. J. Neurosci. 29, 6418–6426. 10.1523/JNEUROSCI.1142-09.2009.

132. Heinricher, M.M. (2004). Principles of Extracellular Single-Unit Recording. In Microelectrode Recording in Movement Disorder Surgery, pp. 8–13. 10.1055/b-0034-56092.

133. Hayat, M., Khan, A., and Yeasin, M. (2012). Prediction of membrane proteins using split amino acid and ensemble classification. Amino Acids. 10.1007/s00726-011-1053-5.

134. Maroco, J., Silva, D., Rodrigues, A., Guerreiro, M., Santana, I., and De Mendonça, A. (2011). Data mining methods in the prediction of Dementia: A real-data comparison of the accuracy, sensitivity and specificity of linear discriminant analysis, logistic regression, neural networks, support vector machines, classification trees and random forests. BMC Res. Notes 4, 299. 10.1186/1756-0500-4-299.

135. Ribeiro, M.T., Singh, S., and Guestrin, C. (2016). “Why should i trust you?” Explaining the predictions of any classifier. In Proceedings of the ACM SIGKDD International Conference on Knowledge Discovery and Data Mining 10.1145/2939672.2939778.

